# NEURAL CORRELATES OF SUBJECTIVE FOOD VALUATION IN THE CONTEXT OF BARIATRIC SURGERY

**DOI:** 10.64898/2026.08.19.745760

**Authors:** Patrick Gagnon, Amélie Lachance, Mélissa Pelletier, Marianne Legault, Sarah-Kim Ross, Sylvain Iceta, Laurent Biertho, François Julien, Catherine Bégin, Alain Dagher, André Tchernof, Yashar Zeighami, Andréanne Michaud

**Affiliations:** Institut universitaire de cardiologie et de pneumologie de Québec - Université Laval, Québec, QC; École de nutrition, Université Laval, Québec, QC; Centre Nutrition, santé et société (NUTRISS), Institut sur la nutrition et les aliments fonctionnels (INAF), Québec, QC; Département de psychiatrie et de neurosciences, Université Laval, Québec, QC; Département de chirurgie générale, Institut universitaire de cardiologie et de pneumologie de Québec - Université Laval, Québec, QC; École de Psychologie, Université Laval, Québec, QC; Montreal Neurological Institute, McGill University, Montréal, QC; Douglas Research Center, McGill University, Montréal, QC

## Abstract

**Objective:** To examine changes in the neural valuation of high- versus low-calorie stimuli following bariatric surgery and determine whether these changes relate to weight loss at 24 months.

**Methods:** Adults undergoing bariatric surgery completed fMRI scans before surgery and at 4, 12 and 24 months post-surgery while performing the Becker-DeGroot-Marschak auction task to assess willingness-to-pay (WTP) for food stimuli. Linear mixed-effect models tested longitudinal changes in WTP-related blood oxygen level-dependent (BOLD) associations and their interactions with total weight loss at 24 months.

**Results:** WTP for high-calorie foods decreased significantly after surgery, WTP of low-calorie foods remained stable. At 4 months, WTP-BOLD associations for high- versus low-calorie stimuli were enhanced within the frontoparietal control network and right lateral orbitofrontal cortex relative to pre surgery. These early postoperative changes did not correlate with 24-month weight loss. Instead, greater 24-month weight loss correlated with both pre-surgical and long-term changes (24 months versus pre-surgery) in WTP-BOLD associations within the precuneus, inferior parietal cortex and visual cortex.

**Conclusion:** Bariatric surgery induces early reductions in the valuation of high-calorie foods, potentially through enhanced cognitive control and aversive processing. However, long-term weight-loss success appears more strongly related to trait-like neural differences in self-referential and attentional processing.

## 1 Introduction

Obesity is a complex, chronic and relapsing disease associated with numerous adverse health outcomes^1^. Modern food environments, characterized by the widespread availability of energy-dense, highly palatable foods, are thought to play a central role by promoting excessive energy intake, weight gain, and dysregulation of appetite and eating behaviors^2^. Despite the recognized importance of energy-dense foods in the development and maintenance of obesity, the neurobehavioral mechanisms that drive their consumption remain poorly understood.

Bariatric surgeries, a group of procedures that alter the gastrointestinal tract and cause significant weight loss, provide a unique opportunity to study these mechanisms of feeding drive and appetite control^3^. In addition to experiencing substantial and sustained weight loss and marked metabolic improvements, individuals who undergo bariatric surgery frequently report pronounced postoperative changes in appetite, food preferences, and the perceived desirability of calorie-dense foods^4,5^. Alterations in appetite-regulating hormones^6,7^ and in eating behaviors such as food disinhibition^8^ have also been reported. Together, these behavioral and physiological shifts suggest that changes in the neural processes underlying food-related decision making may contribute to the long-term metabolic benefits of bariatric surgery^6^. However, value-based decision circuitry underlying food choice has not been investigated directly following bariatric surgery^5^, which this study addresses.

Functional magnetic resonance imaging (fMRI), a neuroimaging technique that allows inference of brain activity, offers a powerful approach for probing these decision circuits^5,9^. However, most neuroimaging studies in the context of bariatric surgery have relied on food-cue reactivity paradigms, in which participants passively view food images^5^. Although these studies have reported changes in reward-related and cognitive control networks^5^, passive cue exposure captures only a limited component of food-reward related processing, which is one of many components of behavior related to food intake, and does not directly probe the neural computations involved in valuation and decision making^10^.

Subjective valuation paradigms provide a more direct approach to studying these processes^10–12^. The Becker-DeGroot-Marschak (BDM) auction task^13^, widely used to assess willingness-to-pay (WTP) for food items, serves as a validated proxy for subjective valuation^14,15^. When performed during fMRI, the task engages neural circuits involved in value computation and decision making, including prefrontal and striatal regions implicated in reward evaluation and cognitive control^14,16^. Although auction paradigms have been used to study food valuation in healthy individuals and in populations with obesity^14,17–23^, they have not yet been used to examine whether and how neural representations of food value change following bariatric surgery.

To address this gap, we conducted a prospective fMRI study to examine neurobehavioral changes during a BDM food auction task before and at 4, 12 and 24 months after bariatric surgery compared to the pre-surgery visit, with a focus on the valuation of high- versus low-calorie stimuli. We further investigated whether neural correlates of subjective valuation were associated with long-term weight-loss outcomes 24 months post-surgery. To our knowledge, this represents one of the largest fMRI studies of bariatric surgery to date. By probing neural activity related to subjective food valuation, this study provides insight into the underlying mechanisms that shape food-related decision making and the valuation of energy-dense foods after bariatric surgery.

## 2 Methods

### 2.1 Participant recruitment

Participants (n=89) were recruited through the elective bariatric surgery program at the *Institut universitaire de cardiologie et de pneumologie de Québec-Université Laval (IUCPQ-UL)* in Québec, Canada. All participants were enrolled in the larger prospective REMISSION study, which investigates determinants of metabolic recovery following bariatric surgery. The cohort consisted of adults with severe obesity undergoing either a sleeve gastrectomy (SG)^24^, a biliopancreatic diversion with duodenal switch (BPD-DS)^25^, or a Roux-en-Y gastric bypass (RYGB)^26^. Surgical techniques have been described previously^27^. Inclusion criteria included being between 18 and 60 years of age and followed NIH guidelines for bariatric surgery: BMI ≥ 40 kg/m², or BMI ≥ 35 kg/m² with obesity-related comorbidities^27,28^. Exclusion criteria included uncontrolled medical, surgical, neurological, or psychiatric disorder, liver cirrhosis or hypoalbuminemia, pregnancy, or history of substance or alcohol abuse^27^. Individuals with prior surgeries involving the stomach, esophagus, brain, or previous bariatric procedures were excluded^27^. Additional exclusions applied to those with gastrointestinal inflammatory conditions or ulcers, severe food allergies, or any contraindication to MRI (implanted medical device, metal fragment in body, or claustrophobia) or to food-related fMRI (use of medications affecting the central nervous system e.g., antipsychotics)^27^. The study protocol was approved by the research ethics committee of the *Centre de recherche de l’IUCPQ-UL* (approval number: 2016–2569). All participants provided written informed consent prior to enrollment.

### 2.2 Study design and experimental procedure

The study design has been previously described by Michaud and colleagues^27^. Assessments were conducted at four timepoints: baseline (approximately two months prior to surgery), and at 4 months, 12 months, and 24 months post-surgery. MRI scans were conducted on the morning of each scheduled visit. To standardize hunger levels, participants fasted for 12 hours prior and consumed a standardized nutritional drink (237 ml Boost Original, Nestlé Health Science; 240 kcal, 41 g carbohydrates, 10 g of protein and 4 g of lipids) 1-2 hours before each MRI session. At each visit, and prior to scanning, individuals completed a comprehensive physical assessment. Body composition was measured using calibrated bioelectrical impedance devices (InBody520, Biospace, Los Angeles, California or Tanita DC-430 U, Arlington Heights, IL). Anthropometric measures (height, waist, hip and neck circumferences) and blood pressure were obtained using standardized protocols^27^. Body mass index (BMI) and total weight loss percentage were calculated according to established protocols^27^. Blood samples were collected at each visit. Appetite-regulating hormones (addition of fasting non- and acylated ghrelin for total ghrelin, postprandial GLP-1 and PYY) were analyzed as previously described^7^. Plasma cholesterol, high-density lipoproteins (HDL), low-density lipoproteins (LDL), triglycerides, glucose, and insulin were measured as part of routine clinical monitoring. The homeostatic model assessment for insulin resistance (HOMA-IR) was calculated as follows: fasting glucose (mmol/L) x fasting insulin (pmol/L) / 135^29^. Hunger levels were assessed before and after the MRI scan using a 10-point visual analog scale. Participants also completed the Three-Factor Eating Questionnaire (TFEQ-51) to assess dietary restraint, disinhibition and susceptibility to hunger^30^, with higher scores indicating greater levels of each behavior.

### 2.3 fMRI paradigm

Participants completed a food valuation task based on the BDM auction paradigm during fMRI acquisition. Food stimuli consisted of pictures of 45 snack items: 15 high-calorie sweet (e.g., chocolate), 15 high-calorie savory (e.g., chips) and 15 low-calorie items (e.g., fruits, vegetables). These items were selected from a bank of 70 food images based on each participant’s familiarity and liking ratings, collected on a 0–5 visual analog scale prior to the pre-surgery scan. Only images rated as familiar were retained, and among these, the most highly liked items were selected for each participant. Stimuli were therefore individualized, selected from the common image bank based on each participant’s own ratings (Fig. 1). Participants also rated perceived caloric density for each item on a 0-5 visual analog scale. Stimuli were created in-house or adapted from a previously published protocol^17^. Detailed stimuli characteristics are described in Supplementary Materials (section 1.1 and Fig. S1-S5).

**Figure 1:**
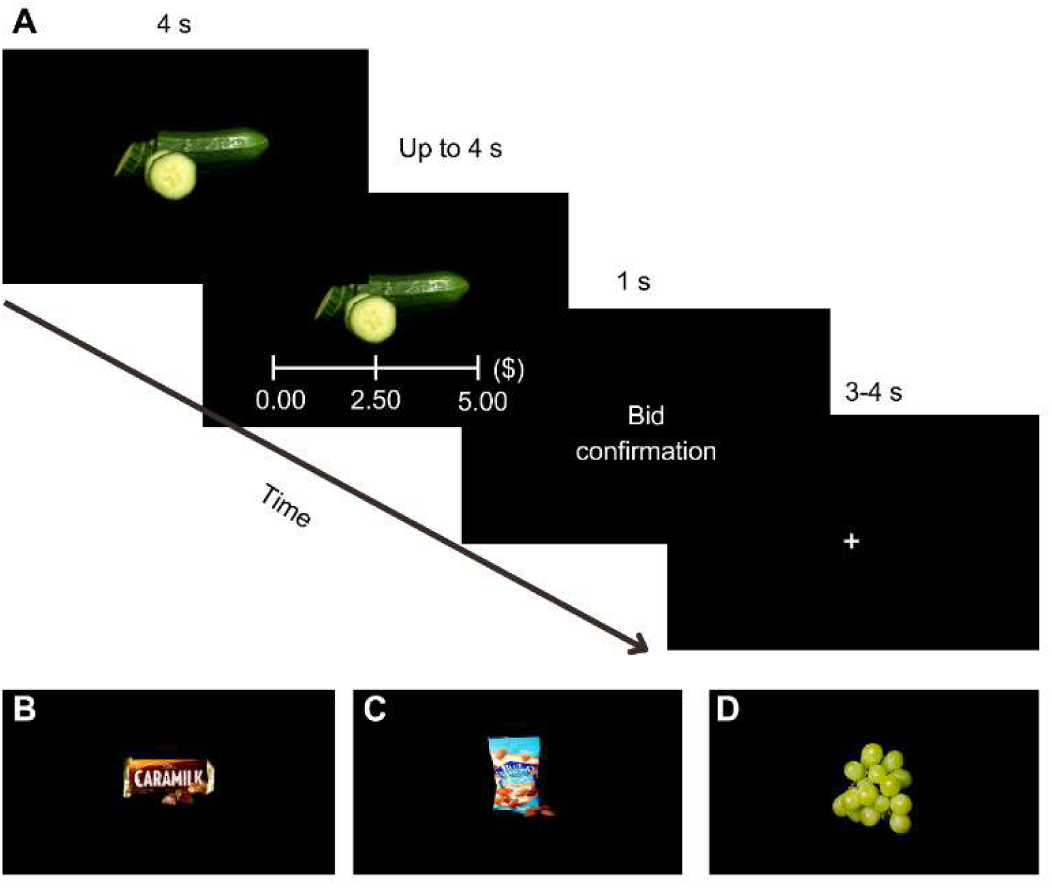
Description of the food auction task. (A) Example of a single trial from the fMRI BDM food auction task. Each trial began with the presentation of a food image, followed by a willingness-to-pay (WTP) screen asking, “How much are you willing to pay for this snack?”. Using an MRI controller, participants indicated their bid on a scale ranging from $0 to $5 in $0.50 increments. The selected bid was then displayed on a confirmation screen (e.g., “Your bid is: $5”). Trials ended with a fixation cross presented for a jittered inter-trial interval. (B) Representative high-calorie sweet food stimulus. (C) Representative high-calorie savory food stimulus. (D) Representative low-calorie food stimulus.

During scanning, participants performed a computerized BDM auction task implemented in E-Prime (Versions 2.0 and 3.0, Psychology Software Tools, Pittsburgh, PA). Visual stimuli were presented via a Philips custom mirror for the head coil reflecting a 32’’ SensaVue television. Each trial began with a 4-second presentation of a food item, followed by a response window of up to 4 seconds during which participants were asked to indicate their WTP (bid amount) between $0 and $5 in $0.50 increments^17^. A fixation cross was then displayed for 3-4 seconds. Each participant completed three runs (∼10 minutes each), during which all 45 items were presented in randomized order.

At the end of the session, one trial was randomly selected for a real auction. A computer-generated price between $0 and $5 determined the outcome: if the participant’s bid exceeded the computer’s price, they received the item and paid the computer-generated amount; otherwise, they kept the full $5 but did not receive the item. Participants were instructed that the optimal strategy was to bid their true WTP^13^.

### 2.4 MRI acquisition

Imaging was performed at the *Centre de recherche de l’IUCPQ-UL* using a 3-Tesla Philips Ingenia whole-body MRI system equipped with a 32-channel head coil. Participants were positioned supine in the scanner.

The imaging protocol included high-resolution T1-weighted 3D turbo field echo images, resting-state fMRI, and task-based fMRI using food-related stimuli. The present study only reports task-based fMRI data. T1-weighted images were acquired with 176 sagittal slices (1 mm isotropic resolution; TR = 8.1 ms; TE = 3.7 ms; FOV= 240 × 240 mm²). The BDM food valuation task was acquired using T2*-weighted echo-planar imaging (EPI) to measure blood-oxygen-level-dependent (BOLD) signals. Each volume included 45 transverse slices (TE = 30 ms, TR = 2750 ms, voxel size 3 × 3 × 3 mm, FOV = 240 × 240 mm², and a flip angle = 80°), for a total task fMRI duration of approximately 30 minutes.

### 2.5 fMRI preprocessing

Preprocessing was performed using fMRIPrep (version 25.1.3)^31^. Briefly, T1-weighted images were corrected for intensity non-uniformity, skull-stripped, segmented into cerebrospinal fluid (CSF), white matter (WM) and grey matter (GM), and normalized to the MNI152NLin2009cAsym template^32^. BOLD images were corrected for head motion and distortions (using field maps), co-registered to the pre-surgery T1-weighted image, and normalized to the same MNI space. Confound regressors including six motion parameters and global CSF and WM signals were extracted for noise correction. A complete fMRIPREP boilerplate is provided in Supplementary Materials (Section 1.2), as recommended by the authors of the software^31^. Quality control procedures, aimed at discarding scans corrupted by excessive head motion, artefacts, or signal loss, are described in Supplementary Materials (Section 1.3).

### 2.6 Statistical analysis

#### 2.6.1 Descriptive statistics

Descriptive anthropometric, cardiometabolic and psychological variables were analyzed using linear mixed-effect models with the Python statsmodels library^33^ to assess longitudinal effects of bariatric surgery across sessions (pre-surgery; 4, 12, 24 months post-surgery). For each model, uncorrected p-values from Wald test chi-square tests for the session effect were reported^33^.

#### 2.6.2 Task behavioral analysis

For each participant and session, WTP values from the BDM task were averaged separately for high- and low-calorie stimuli. These values were analyzed using a mixed-effects model with session (pre-surgery, 4, 12 and 24 months post-surgery) and calorie density (high vs. low) as categorical variables, along with their interaction. Covariates included surgery type (SG, RYGB, BPD-DS), sex, baseline BMI, and baseline age. Participant was modeled as a random effect. Exploratory mixed effect models were also computed to quantify the associations between WTP for high-calorie stimuli and appetite-regulating hormones as well as TFEQ scores, with Bonferroni correction applied. Correlations were computed between WTP and numerous stimuli properties to replicate former findings (true caloric density, estimated caloric density, liking and retail price)^17–19^ as well as for stimuli visual confounds differing significantly between high and low-calorie stimuli^34^.

#### 2.6.3 fMRI – participant-level modeling

Voxel-wise participant-level general linear models were estimated using Nilearn^35^. To model BOLD responses to food cues, three boxcar regressors corresponding to viewing periods of high-calorie sweet, high-calorie savory, and low-calorie items were convolved with the Glover hemodynamic response function. Subjective valuation was modeled using three additional regressors for the same viewing periods, with amplitudes modulated by run mean-centered WTP values (reflecting the association between WTP and BOLD percent signal change, i.e. parametric modulation). Two contrasts were computed: (1) parametric modulation while viewing high- versus low-calorie stimuli and (2) unmodulated viewing of high- versus low-calorie stimuli. For visualisation, parametric modulation contrasts while viewing high- and low-calorie stimuli were computed separately. This procedure was applied across all participants, sessions, and runs, yielding statistical β contrast maps of BOLD percent signal change per dollar of WTP (modulated contrasts) or unmodulated percent signal change, which were used for group-level analyses. Additional details are provided in Supplementary Materials (Section 1.4).

#### 2.6.4 fMRI – Group-level modeling

Participant-level contrast maps across the runs were entered into voxel-wise linear mixed-effects models using AFNI - 3dLMEr^36^ to assess longitudinal effects of bariatric surgery (pre-surgery; 4, 12, 24 months post-surgery) for both unmodulated and parametric modulation contrasts.

**Model I** (Session effect):

Voxel β_contrast_ = Session + Surgery type + Handedness + Sex + Run + BMI_baseline_ + Age_baseline_ + Hunger_pre-scan_ + 1 | participant

A second model examined whether WTP–BOLD percent signal change associations interacted with total weight loss at 24 months, restricted to participants with available total weight loss data.

**Model II** (Session x Total Weight Loss):

Voxel β_contrast_ = Session x TotalWeightLoss_24months_ + Surgery type + Handedness + Sex + Run + BMI_baseline_ + Age_baseline_ + Hunger_pre-scan_ + 1 | participant

Cluster-level correction was performed using AFNI’s 3dClustSim with 10,000 iterations, using a voxel-wise threshold of p < 0.001 and a family-wise error (FWE) rate of α = 0.05 (two-sided)^37^, resulting in a minimum cluster size of 33 voxels (NN1). Thresholded group-level maps included session means, session contrasts (e.g., 4 months vs. pre-surgery, 12 months vs. pre-surgery, and 24 months vs. pre-surgery), and model covariates. Significant clusters were characterized by computing spatial overlap with Brainnetome atlas regions^38^, expressed as the percentage of cluster volume overlapping each region^39^. Additional details are provided in the Supplementary Materials (Section 1.5).

To examine surgery-related changes in functional networks involved in food valuation, participant-level maps for the modulated contrast (high- versus low-calorie) were parcellated into Yeo’s seven canonical functional networks^40^. Mean β_contrast_ coefficients were extracted from each network and analyzed using linear mixed-effects models mirroring Model I. Bonferroni multiple-comparison correction was applied across the seven networks.

## 3 Results

### 3.1 Participant characteristics

Table 1 shows the clinical characteristics of the participants. A total of 57 participants, predominantly women, were included in the study (Flowchart of study participants – Fig. S6). At baseline, participants exhibited metabolic impairments, including insulin resistance and dyslipidemia. The most frequently performed surgery was SG (47%). Participants exhibited substantial weight loss, reaching a mean total weight loss of 22.4% ± 4.2% at 4 months (BMI: 34.0 ± 3.3 kg/m^2^) and 35.4% ± 7.1% at 12 months (BMI: 28.5 ± 3.5 kg/m^2^), which was maintained at 24 months.

**Table 1:**
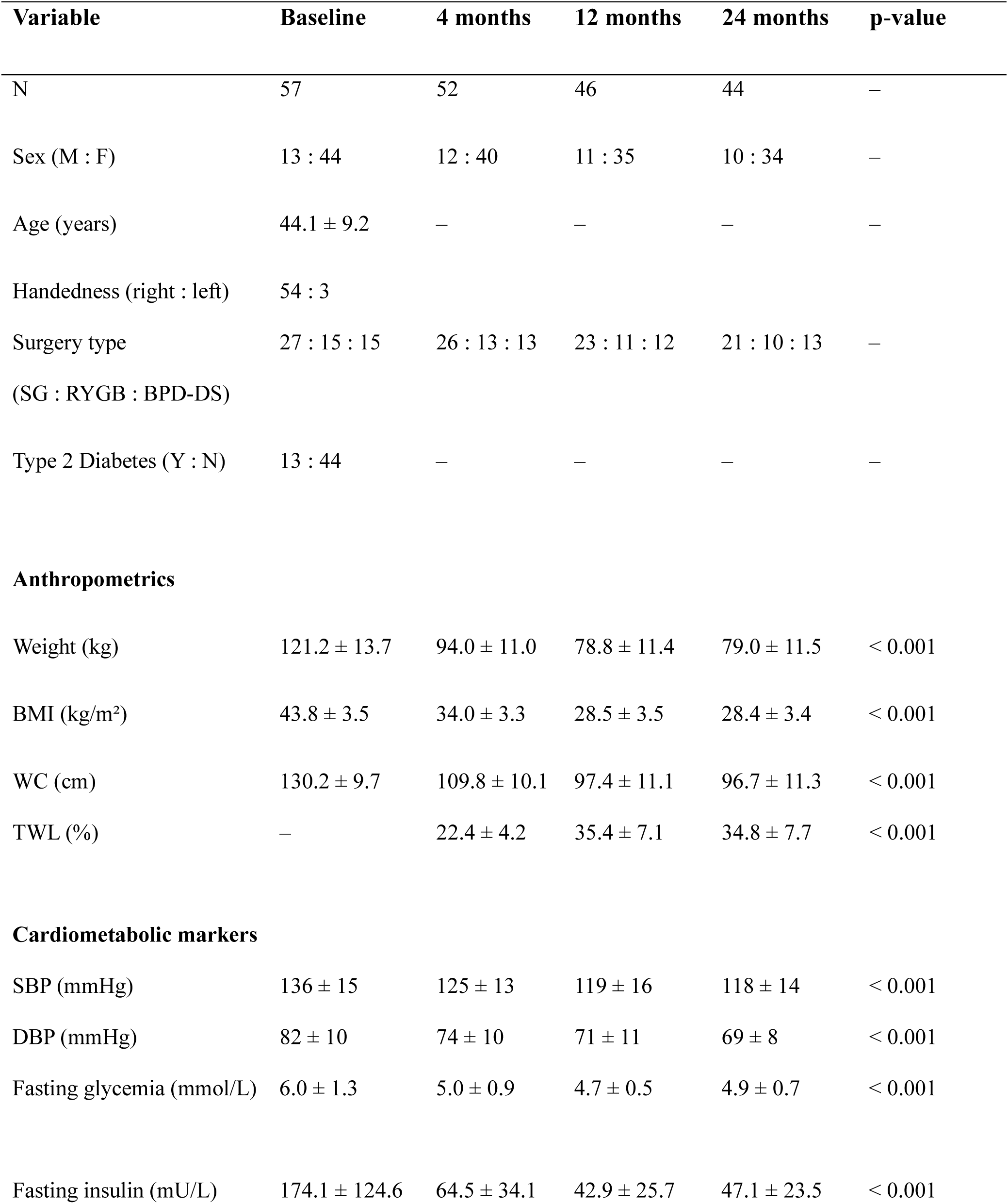
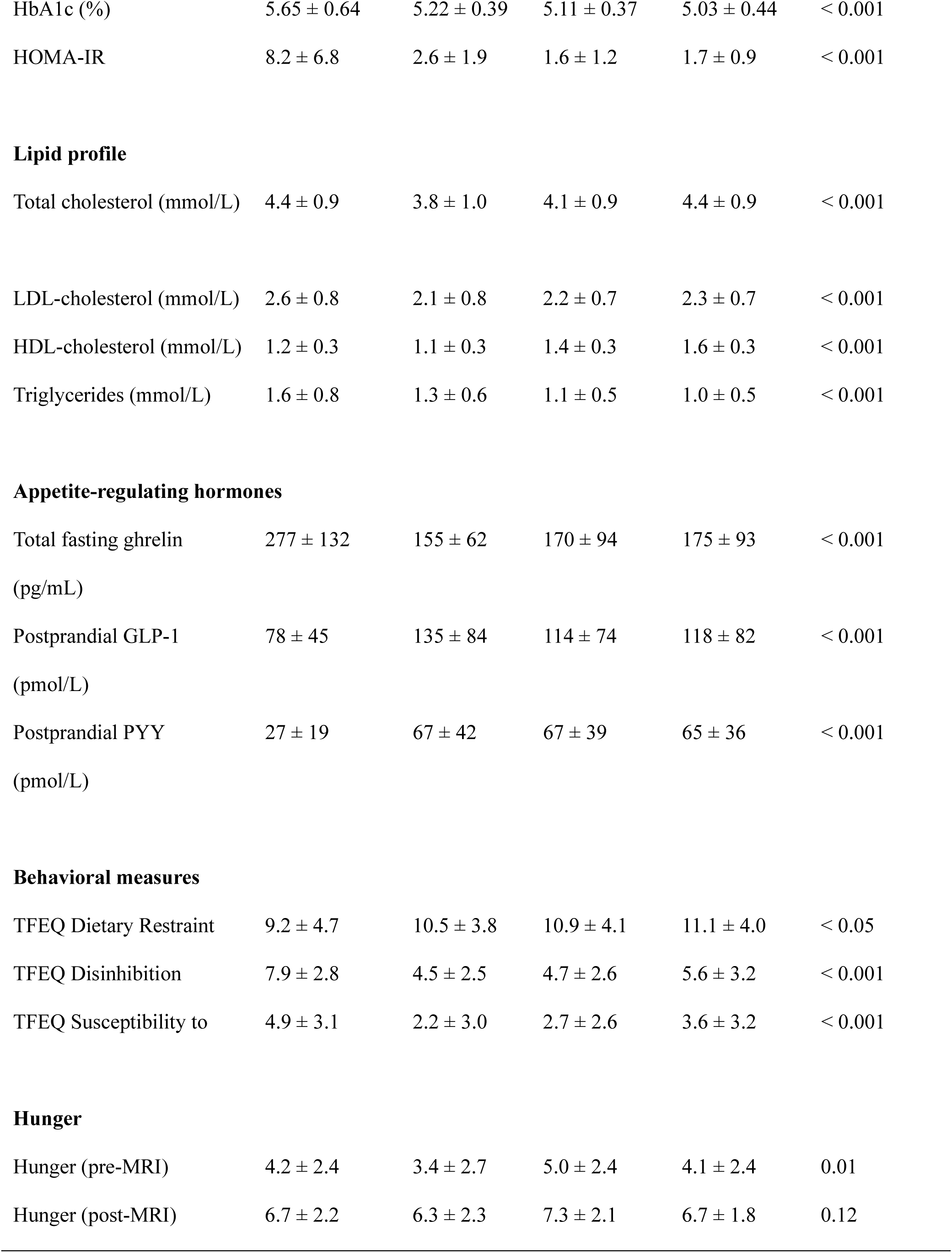
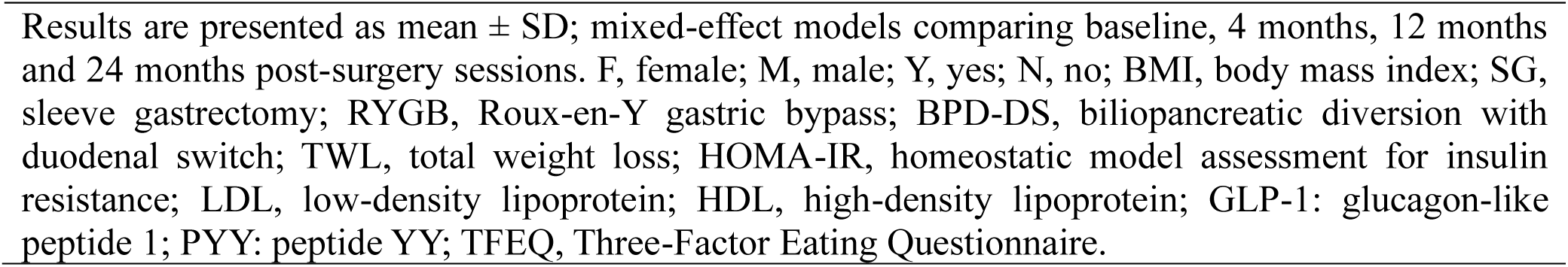
Participant characteristics at baseline and at 4, 12, and 24 months following bariatric surgery.

### 3.2 Cardiometabolic and cognitive outcomes

This was accompanied by significant improvements in cardiometabolic markers, including blood pressure, insulin resistance, and triglyceride levels, as well as significant changes in appetite-regulating hormones, including decreased fasting ghrelin levels and increased postprandial GLP-1 and PYY levels. After surgery, eating behavior scores on the TFEQ shifted toward higher cognitive restraint and lower disinhibition and susceptibility to hunger, and remained significantly different from baseline throughout the study.

### 3.3 Reduced willingness-to-pay for high-calorie stimuli following bariatric surgery

At baseline (i.e., pre-surgery), WTP did not differ between high-calorie ($2.41 ± 0.74) and low-calorie stimuli ($2.45 ± 0.80, p=0.72; Fig. 2A). Following surgery, WTP for high-calorie stimuli decreased significantly at 4 months ($1.65 ± 0.89, p<0.001) and remained significantly lower than baseline at 12 months ($1.96 ± 0.76, p=0.001) and 24 months ($2.09 ± 0.78, p=0.010). In contrast, WTP for low-calorie stimuli did not significantly change across timepoints (4 months: $2.62 ± 0.77, p=0.20; 12 months: $2.65 ± 0.84, p=0.15; 24 months: $2.55 ± 0.81, p=0.79, Fig. 2A).

**Figure 2:**
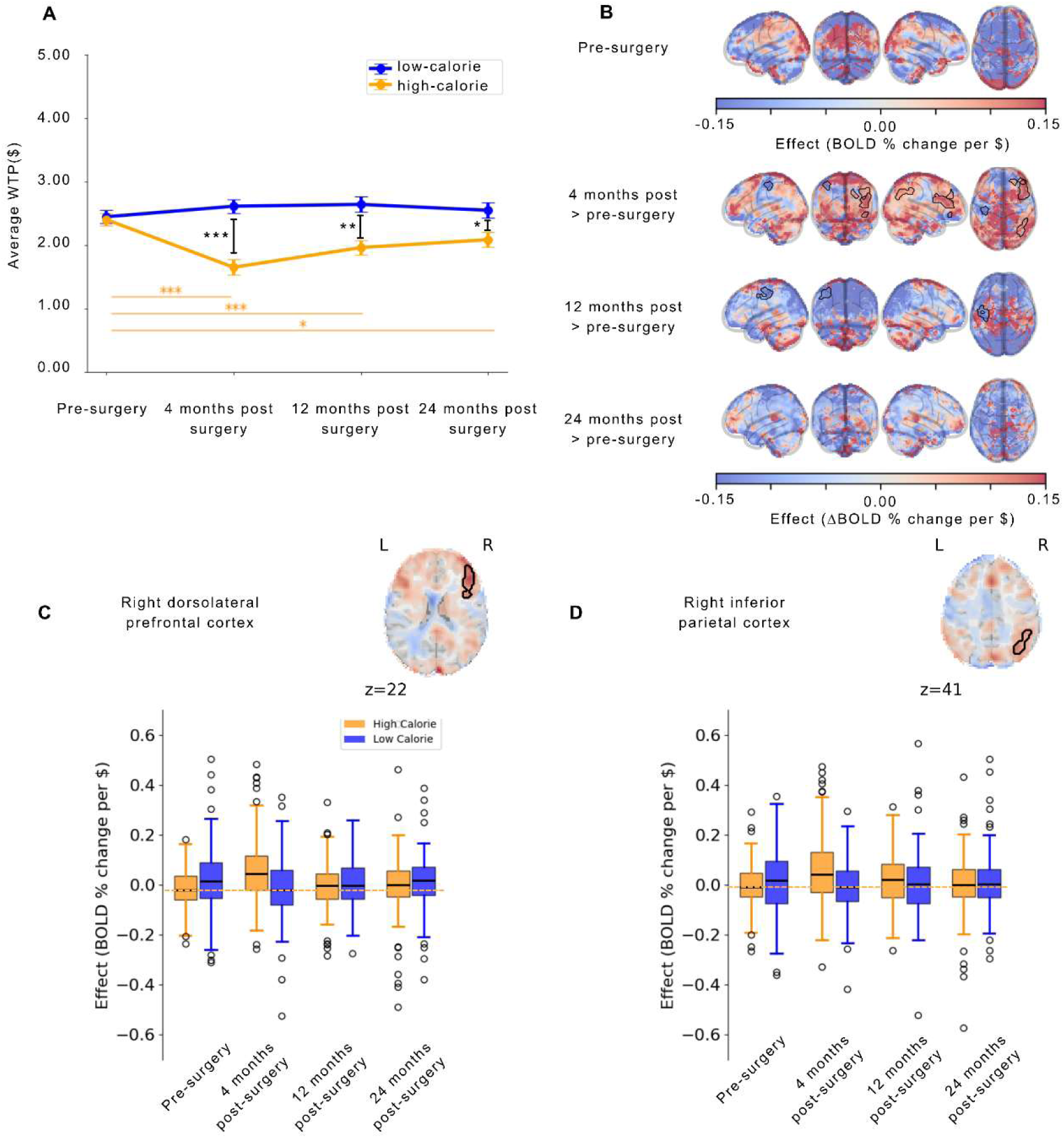
Neurobehavioral correlates of food subjective valuation. (A) Results from the linear mixed-effects model showing willingness-to-pay (WTP) for low- and high-calorie food stimuli across surgical timepoints, adjusted for sex, age, baseline BMI, and surgery type (*p < 0.05; **p < 0.01; ***p < 0.001). Error bars represent the standard error of the mean. (B) Brain regions in which WTP-BOLD percent signal change associations significantly differed for high- versus low-calorie stimuli at pre-surgery or across postsurgical timepoints relative to pre-surgery. Black contours delineate significant clusters for the labeled session contrast (voxel-wise threshold of p < 0.001; cluster-level FWE correction at 5%). The coolwarm colormap represents effect size estimates (BOLD percent signal change per $1 change in WTP relative to the run mean at pre-surgery; or change in BOLD percent signal change per $1 change in WTP relative to the run mean for between-session contrasts). (C) Participant-level mean BOLD percent signal change per dollar of WTP from the largest significant cluster (right dorsolateral prefrontal cortex). (D) Participant-level mean BOLD percent signal change per dollar of WTP extracted from the second-largest significant cluster (right inferior parietal cortex). In panels C and D, boxplots are shown separately for high- and low-calorie stimuli across surgical timepoints, and the yellow dotted line indicates the presurgical BOLD percent signal change per dollar of WTP for high-calorie stimuli.

### 3.4 Associations between willingness-to-pay and BOLD signal change amplitude

#### 3.4.1 Neural correlates of food valuation (Model I)

Prior to surgery, no significant differences were observed in the association between BOLD percent signal change and WTP for high- versus low-calorie stimuli (Fig. 2B).

At 4 months post-surgery compared with pre-surgery, we observed significantly increased associations between WTP and BOLD percent signal change for high- versus low-calorie stimuli in the dorsolateral prefrontal cortex, rostro-dorsal inferior parietal cortex, and lateral orbital gyrus (OFC) (Fig. 2B, Table 2). In contrast, decreased associations were observed in the left postcentral gyrus (Fig. 2B, Table 2). To further characterize these effects, we examined high- and low-calorie stimuli separately by visualizing participant-level session effect sizes within the two primary clusters showing changes at 4 months (Fig. 2C–D). In the dorsolateral prefrontal cortex, WTP-BOLD associations decreased for low-calorie stimuli but increased for high-calorie stimuli (Fig. 2C). A similar pattern was observed in the inferior parietal cortex (Fig. 2D).

**Table 2:**
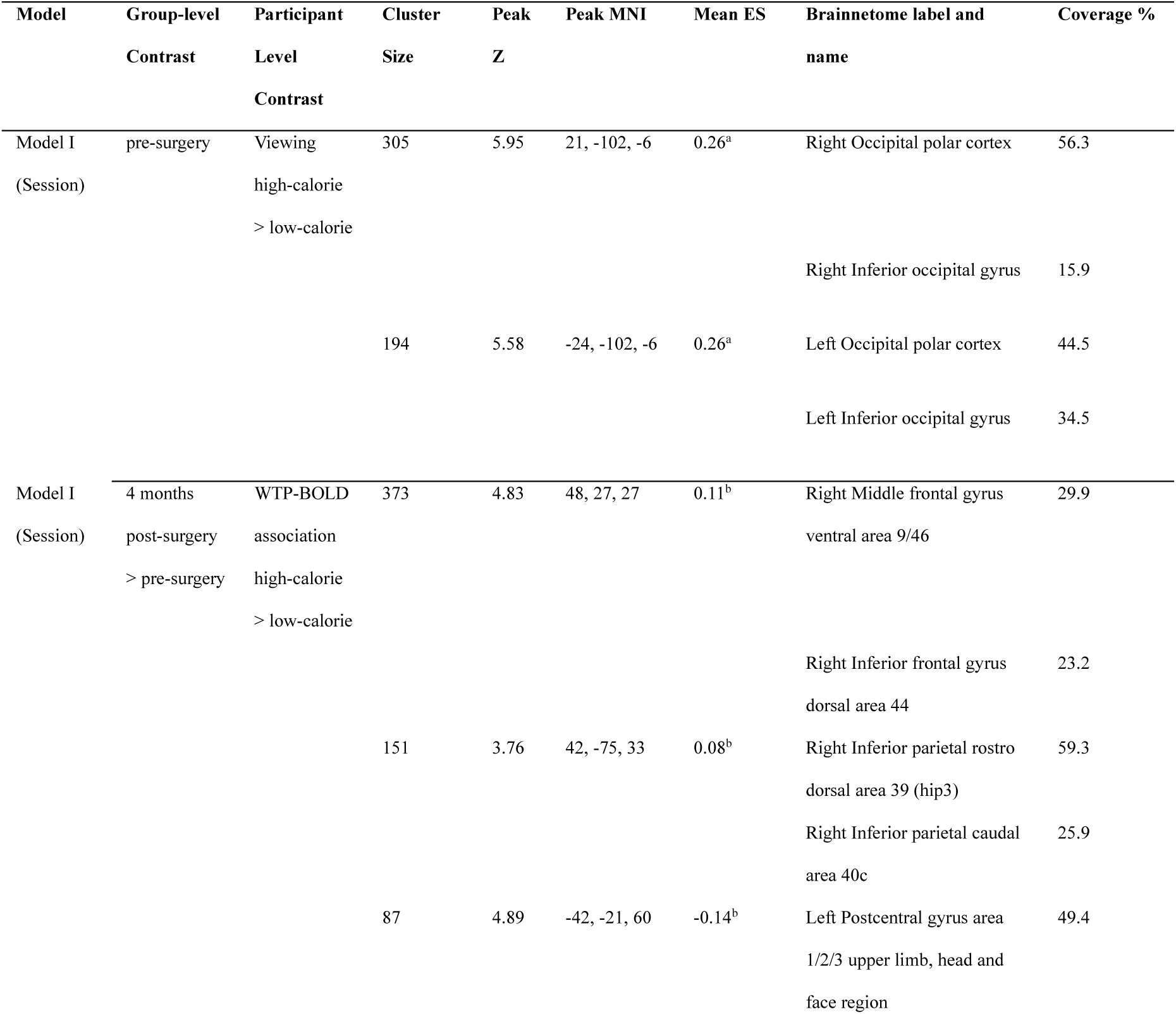
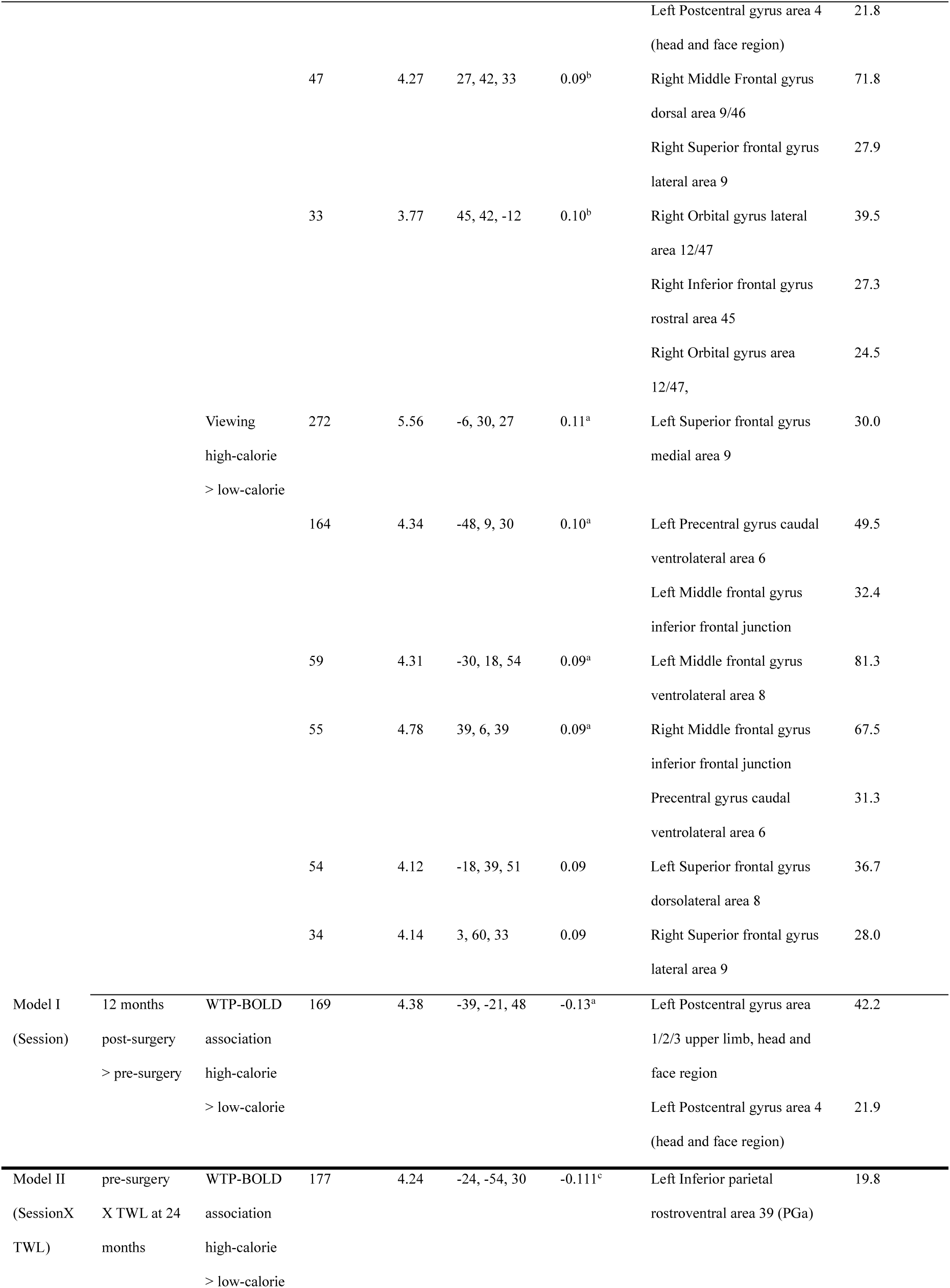
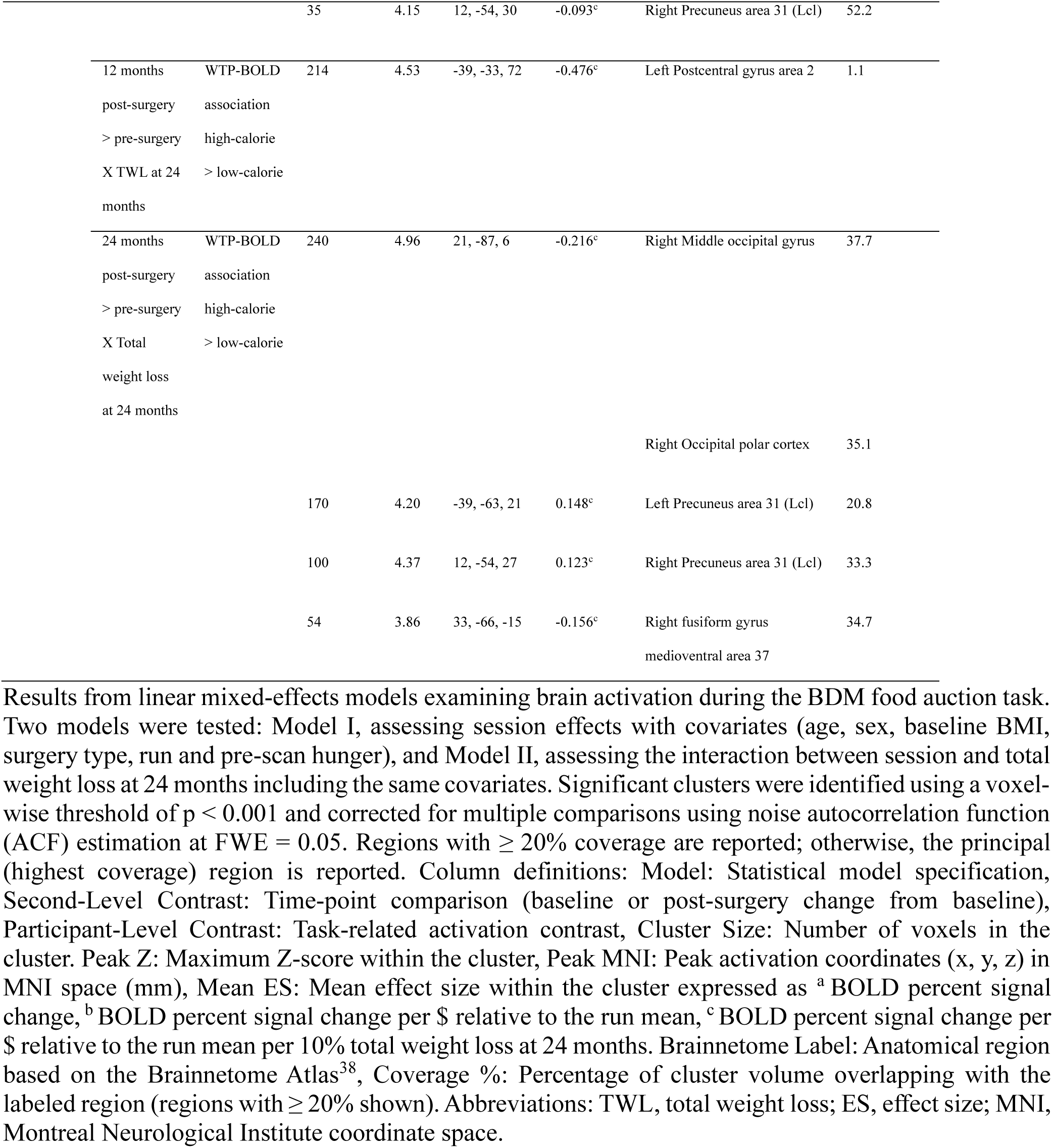
Significant clusters identified in the whole-brain analyses for Models I and II.

At 12 and 24 months post-surgery, minimal changes in WTP–BOLD associations were observed compared to pre-surgery (Fig. 2B). At 12 months, a significant decrease in WTP–BOLD association for high- versus low-calorie stimuli was observed in the postcentral gyrus, whereas no significant changes were observed at 24 months (Fig. 2B). Clusters associated with covariates are reported in Table S1.

Unmodulated responses during food viewing revealed higher BOLD percent signal change for high-calorie compared with low-calorie stimuli, mainly in the visual cortex (Fig. 3, Table 2), reflecting enhanced visual processing of energy-dense foods. Changes in unmodulated BOLD percent signal change between high- and low-calorie stimuli were also evident at 4 months post-surgery as compared to pre-surgery, with increased signal in the superior frontal gyrus, precentral gyrus and middle frontal gyrus (Fig. 3, Table 2).

**Figure 3:**
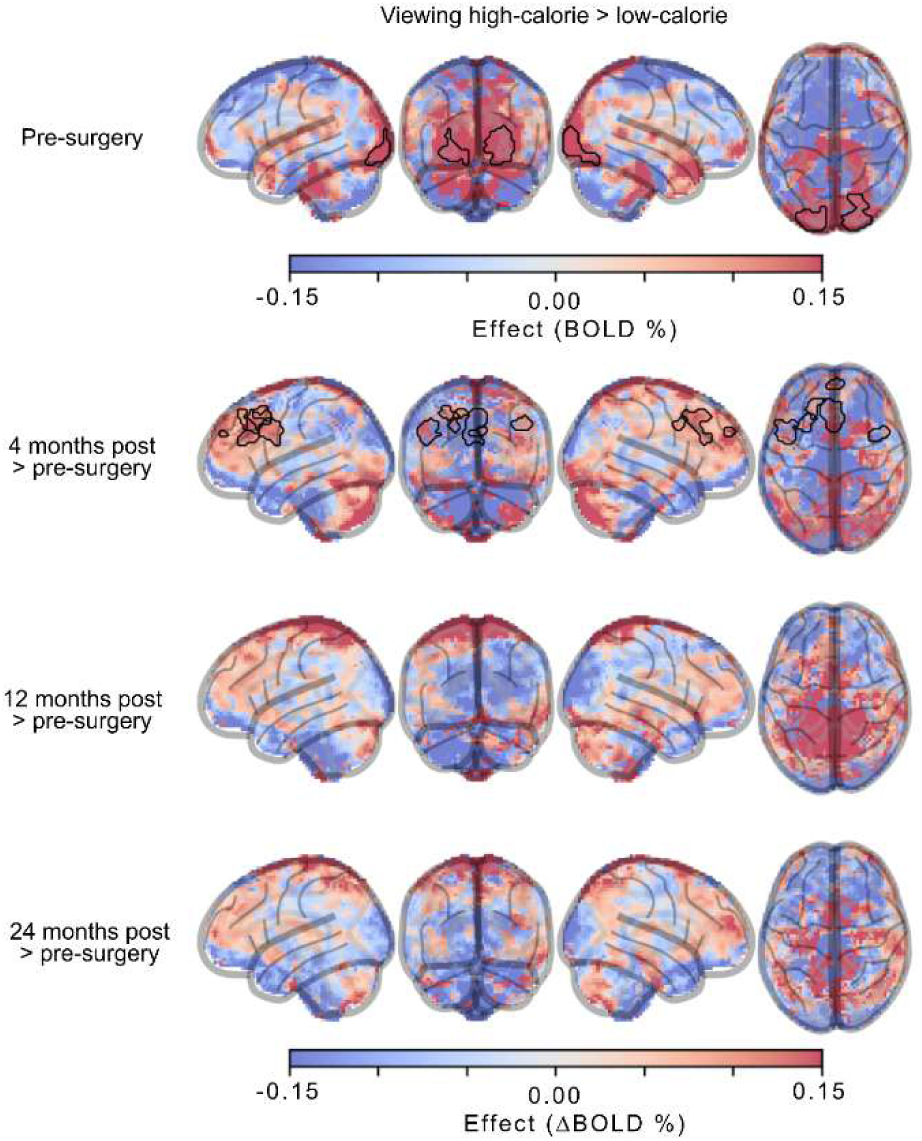
Brain regions in which unmodulated BOLD percent signal change significantly differed between high- versus low-calorie stimuli at pre-surgery or across postsurgical timepoints relative to pre-surgery. Black contours delineate significant clusters (voxel-wise threshold of p < 0.001; cluster-level FWE correction at 5%). The coolwarm colormap represents effect size estimates (BOLD percent signal change at pre-surgery or change in BOLD percent signal change for between-session contrasts).

To better understand the variability in WTP for high-calorie stimuli, we examined associations with TFEQ subscales (disinhibition, susceptibility to hunger, cognitive restraint), and circulating levels of fasting total ghrelin and postprandial PYY and GLP-1 (Fig. 4). WTP for high-calorie stimuli was significantly and positively associated with disinhibition, susceptibility to hunger, and fasting total ghrelin (Fig. 4). Associations between WTP and food stimulus characteristics are shown in Fig. S7-S14 and detailed in Supplementary Materials (Section 2.1).

**Figure 4:**
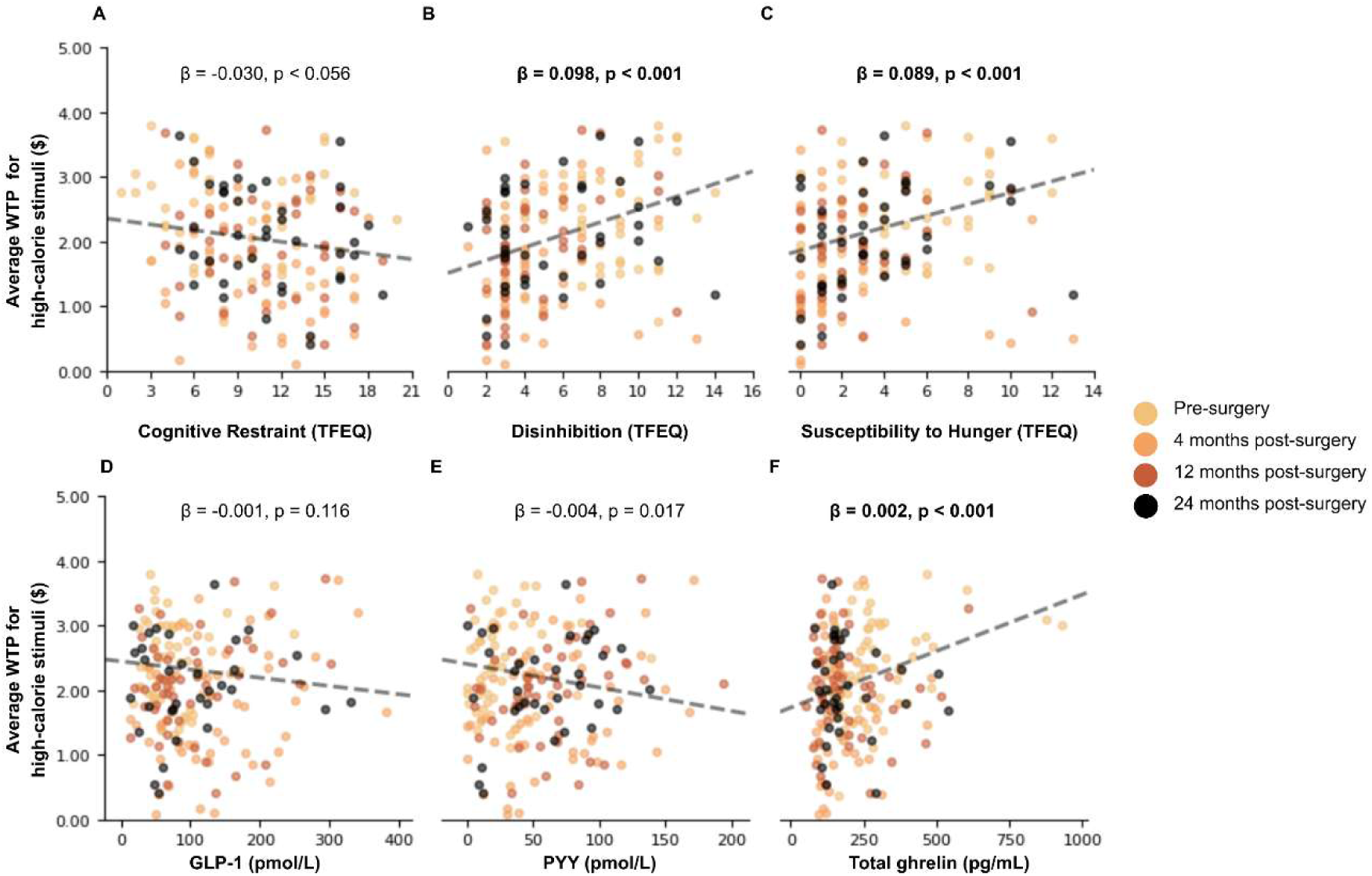
Longitudinal associations between willingness-to-pay (WTP) for high-calorie stimuli and gastro-intestinal hormones or eating behaviours. Associations for (A) cognitive restraint, (B) disinhibition, (C) susceptibility to hunger (TFEQ subscales), (D) postprandial glucagon-like peptide-1, (E) postprandial peptide YY, and (F) fasting total ghrelin. Scatterplots show all observations across sessions; colors denote individual sessions and are included for visualization purposes only. The fitted line represents the beta coefficient from a linear mixed-effects model estimating the association between each predictor (x-axis) and WTP for high-calorie stimuli across all timepoints. Models were adjusted for surgery type, sex, age, and baseline BMI. Statistical significance was assessed using a Bonferroni correction for six comparisons; bold text indicates results that remain significant after correction.

#### 3.4.2 Functional network parcellation analysis (Model I)

When examining WTP-BOLD associations for high- versus low-calorie stimuli within Yeo’s seven functional networks, only the frontoparietal control network showed a significant increase in WTP-BOLD association at 4 months post-surgery compared with pre-surgery (p < 0.05, Bonferroni corrected, Fig. 5), indicating engagement of cognitive control processes during food valuation early after surgery.

**Figure 5:**
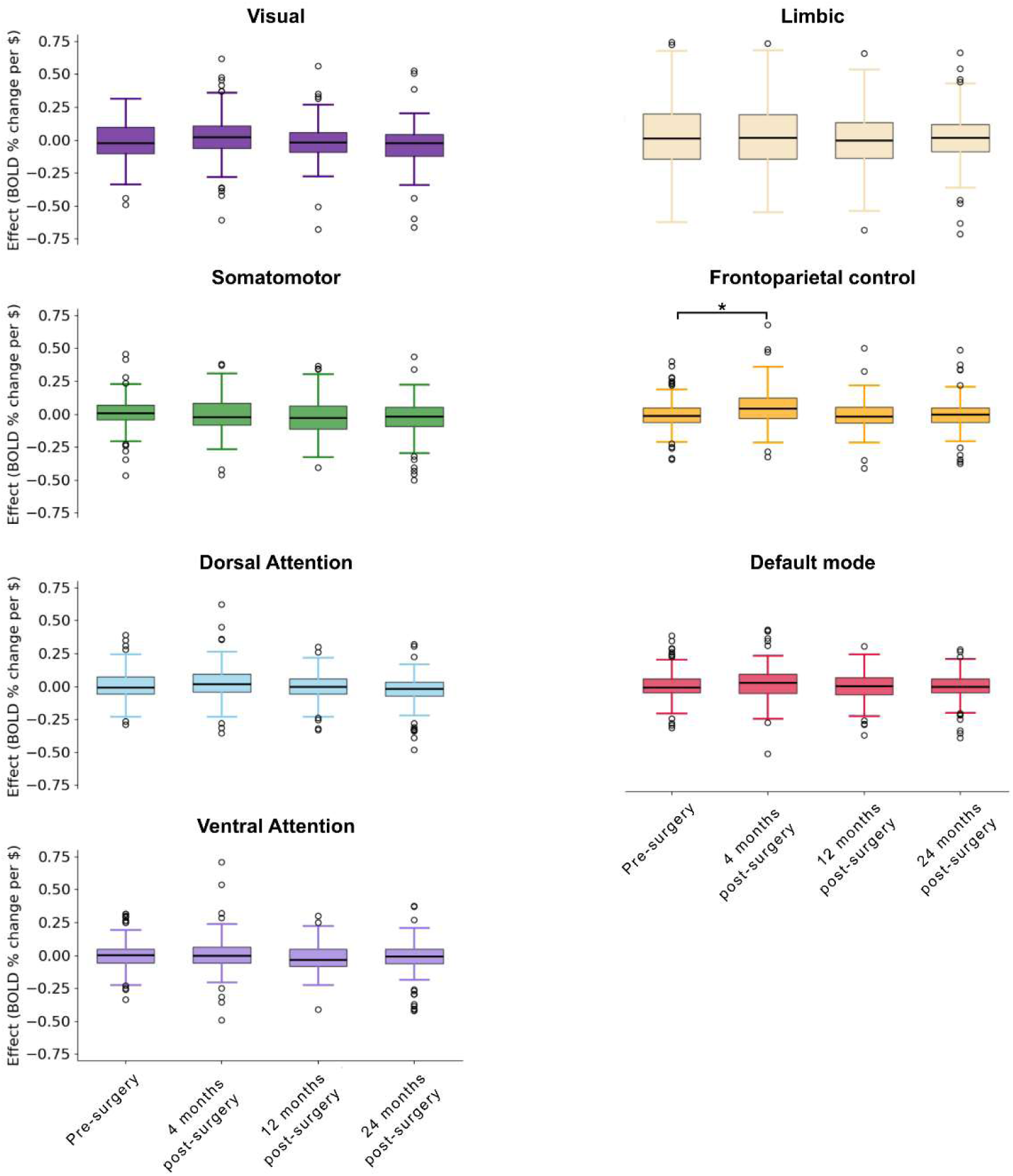
Boxplots showing the distribution of WTP-BOLD association coefficients for high-calorie > low-calorie contrast across the seven Yeo functional networks. Coefficients were extracted from participant-level general linear models. Results from the network-wise mixed-effects model that remained significant after Bonferroni correction for seven tests are indicated with an asterisk (*).

#### 3.4.3 Neural correlates of subjective valuation and their interaction with total weight loss at 24 months (Model II)

To determine whether neural valuation signals were associated with long-term weight loss, we examined the interaction between WTP–BOLD associations for high- versus low-calorie stimuli and total weight loss at 24 months post-surgery (n = 51). At pre-surgery, two clusters in the right precuneus and left inferior parietal cortex showed significant associations with total weight loss at 24 months (Fig. 6A). In both clusters, higher WTP–BOLD associations for high- versus low-calorie stimuli were associated with lower total weight loss at 24 months (Fig. 6B-C). No significant clusters were identified for changes in WTP-BOLD associations at 4 months post-surgery relative to pre-surgery. At 12 months post-surgery versus pre-surgery, a cluster in the postcentral gyrus showed a significant interaction with total weight loss at 24 months. At 24 months post-surgery relative to pre-surgery, WTP-BOLD associations showed a negative interaction with total weight loss in the occipital gyrus and fusiform gyrus, and a positive interaction in bilateral precuneus, overlapping visually with pre-surgery clusters (Fig. 6A). Clusters associated with covariates are reported in Table S1.

**Figure 6:**
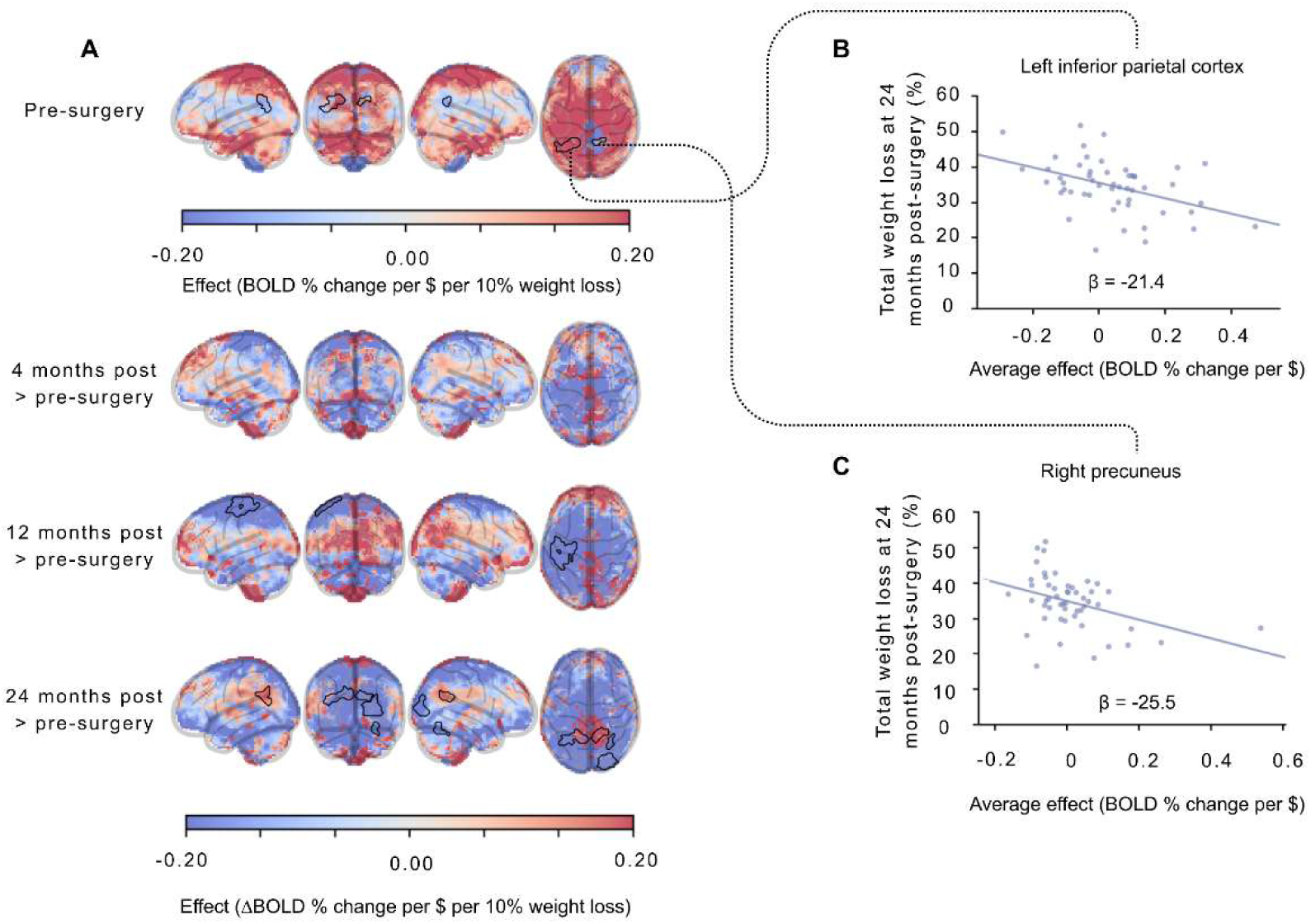
(A) Brain regions where WTP-BOLD percent signal change associations for high- versus low-calorie stimuli showed a significant interaction between session (pre-surgery, 4, 12 and 24 months) and total weight loss and 24 months. Black contours delineate significant clusters for the displayed session contrast (voxel-wise threshold p < 0.001; cluster-level correction at 5% FWE). The coolwarm colormap represents effect size estimates (BOLD percent signal change per WTP dollar relative to the run mean, per 10% total weight loss for pre-surgical effects; or change in BOLD percent signal change per WTP dollar relative to the run mean, per 10% total weight loss for between-session contrasts). For visualization, scatterplots of mean percent signal change per WTP dollar and total weight loss at 24 months are shown for two significant clusters identified at pre-surgery in (B) left inferior parietal cortex and (C) right precuneus. β denotes the regression coefficient of the fitted line, representing the change in % total weight loss per unit of WTP-related BOLD percent signal change/$. Models were adjusted for baseline age, BMI, sex, surgery type, handedness and hunger levels.

## 4 Discussion

Our study provides the first evidence suggesting that bariatric surgery reduces the valuation of high-calorie food stimuli, with limited impact on low-calorie stimuli. These early changes were associated with increased engagement of frontoparietal cognitive control networks and lateral orbitofrontal cortex 4 months after surgery. Together, these findings support a model in which bariatric surgery induces an early, and partly transient, reorganisation of food valuation processes. Importantly, these early postoperative neural changes did not correlate with long-term weight loss, suggesting that they may reflect state-dependent adaptations. In contrast, long-term weight loss was associated with pre-surgical and long-term postoperative neural activity in the precuneus, inferior parietal cortex and visual cortex, regions involved in self-referential^41^ and attentional^42^ processing. These findings suggest that individual differences in how food cues are perceived and integrated into valuation processes may play a more central role in long-term surgical outcomes than cognitive control mechanisms, which are limited to the early postoperative period.

Although WTP has not previously been examined in the context of bariatric surgery, our findings align with a broad literature showing reduced wanting, craving, and in some cases liking for high-calorie foods following surgery^5^. Across all timepoints, WTP for high-calorie stimuli was positively associated with susceptibility to hunger and disinhibition (TFEQ scores), as well as circulating ghrelin levels, all of which typically decrease after bariatric surgery^8,43^. Given ghrelin’s role in enhancing incentive salience of food^44^, these associations are consistent with a reduction in motivational value of high-calorie foods following surgery. Although WTP for high-calorie stimuli remained positively associated with true caloric density after surgery, its association with baseline estimated caloric density became negative after surgery, particularly at 4 months. This suggests a shift in how energy-dense foods are subjectively valued.

Furthermore, the association between pre-surgery liking ratings and WTP for high-calorie stimuli was selectively attenuated at 4 months and re-emerged at 24 months, suggesting a transient decoupling between pre-surgery hedonic appreciation and motivational valuation. However, as liking was not reassessed concurrently at these timepoints, we cannot rule out that this pattern partly reflects changes in liking itself over time.

Our fMRI results further suggest that this early reduction in subjective value assigned to high-calorie stimuli involves greater engagement of cognitive control networks. Concurrently, we found increased WTP-BOLD associations for high- versus low-calorie stimuli in the lateral orbitofrontal cortex (area 12/47), a region implicated in aversion processing, unpleasantness, or in not receiving expected reward, also known as “non-reward” signaling^45^. These findings suggest a reorganisation of value computation, whereby high-calorie foods are processed through enhanced regulatory and affective control systems. Although bariatric surgery is a physiological intervention that alters metabolism, gastrointestinal hormones, and gut-brain signaling, particularly four months post-surgery, it also imposes behavioral changes^46^. Patients receive nutritional counselling, must adhere to dietary guidelines, and often experience gastrointestinal symptoms that limit overeating, especially at the early stage^46^. Together, these factors promote a restrictive eating context, particularly during the early postoperative period, which may require or facilitate increased cognitive control over food valuation in order to avoid excess food intake consequences^47^.

Importantly, these early changes in WTP-BOLD associations were not sustained over time. At 12 and 24 months, we observed minimal changes in WTP-BOLD associations compared with pre-surgery, alongside a gradual return of WTP for high-calorie stimuli toward pre-surgery levels. This pattern suggests that early postoperative alterations in valuation-related and regulatory processes may be transient, potentially reflecting an adaptive phase that evolves as individuals adjust to the post-surgical state, and it may contribute to inter-individual variability in long-term outcomes, including weight regain in a subset of patients^46^.

Our findings are consistent with prior fMRI studies of food cue reactivity and resting-state activity after bariatric surgery^48–50^, which have reported increased engagement of cognitive control networks, although findings are heterogeneous across studies^5^. This variability likely reflects methodological differences between passive cue-reactivity paradigms and the incentive-based valuation process captured by the BDM task, as well as smaller sample sizes in other studies. Also consistent with the literature, baseline results from the unmodulated model replicated prior findings in individuals living with overweight or obesity, including stronger engagement of visual processing regions in response to high- versus low-calorie stimuli^20,42,51^.

fMRI activation patterns correlated with long-term weight loss outcomes. Specifically, WTP-BOLD associations in the precuneus and inferior parietal cortex interacted with total weight loss at 24 months, such that pre-surgical clusters showed a negative relationship, while 24-month postoperative changes within similarly localized clusters showed a positive relationship. In contrast, early postoperative changes (4 months versus pre-surgery) did not significantly correlate with the amount of long-term 24-month weight loss. These findings suggest that baseline and long-term neural signatures, rather than early postoperative changes, are more closely associated with sustained weight loss. Altered involvement of the posterior cingulate and precuneus during food cue reactivity has previously been shown in obesity^41^, highlighting the relevance of self-referential processing in long-term surgical outcomes. In addition, we found that long-term neural changes in the fusiform gyrus and occipital cortex were also significantly associated with weight loss at 24 months. Overall, these results may support the notion that obesity is a chronic, relapsing condition^1^. Regardless of how the intervention transiently alters brain activity patterns in the shorter term, long-term outcomes were more strongly associated with pre-surgical brain activity traits. Moreover, partially overlapping regions exhibited inverse associations at baseline and positive associations with 24-month postoperative changes, suggesting a relapse-like reversion toward the pre-surgical state.

Our study is not without limitations. The absence of a healthy control group or an alternative obesity treatment intervention group limits our ability to disentangle effects specific to time, repeated exposure, baseline obesity or the surgical intervention itself. Some potentially relevant individual-level variables (e.g., income, education, dietary habits) were not measured and could have influenced WTP values. However, the longitudinal within-subject design mitigates this limitation by using each participant as their own control. Menstrual cycle phase was not recorded, which may influence food-related neural responses^9^. Despite these limitations, the longitudinal design, relatively large sample size for this field, and the novelty of the BDM-based fMRI paradigm used in the context of bariatric surgery strengthen the study.

## 5 Conclusion

In conclusion, bariatric surgery leads to a reduced subjective valuation of high-calorie stimuli, with minimal effects on low-calorie stimuli. These behavioral changes are related to increased engagement of cognitive control and aversive valuation networks during the early postoperative period. However, these early neural changes are not associated with weight loss outcomes. Instead, both preoperative neural activity and long-term postoperative changes in brain regions involved in self-referential processing and visual processing are associated with long-term weight loss. Further studies are required to examine whether subjective food valuation and its neural correlates can serve as predictors of long-term metabolic outcomes following bariatric surgery or other obesity-related interventions.

## 6. Acknowledgements

We recognize the contributions of surgeons, nurses, and medical staff of the bariatric surgery program at IUCPQ-UL. We thank Catherine Lemieux, undergraduate intern, who helped with the quality control of fMRI scans. We also thank the MRI technicians, Xavier Moreel (Coordinator of the *Plateforme d’imagerie avancée* at IUCPQ-UL), and Guillaume Gilbert (*Philips)* for their support. We thank Lucie Bouffard from Dr. André Carpentier’s laboratory for gut hormone measurements and Mélanie Nadeau for her involvement in the REMISSION study. We also wish to acknowledge the late Denis Richard, a valued collaborator of this study. Special thanks to Zaki Alasmar and Aliza Brzezinski at the Cerebral Imaging Center of the Douglas Institute for their help with MRI preprocessing and analysis. The authors wish to acknowledge the invaluable contribution of the late Justine Daoust, who participated in data analysis and passed away before the completion of this work. We are deeply grateful for her unyielding dedication to this research. Finally, we sincerely thank all the participants who took part in this study.

AI tools assisted with drafting and analysis scripts. All outputs were reviewed and edited by the authors, who take full responsibility.

