## Supplementary material for "NEURAL CORRELATES OF SUBJECTIVE FOOD VALUATION IN THE CONTEXT OF BARIATRIC SURGERY"

### 1 Supplementary notes on methods

#### 1.1 Supplementary analysis of willingness-to-pay (WTP) and stimulus properties

True calorie density and total calories per stimulus were extracted from the Canadian Nutrient File[1] or, when unavailable, from local grocery store websites (e.g., maxi.ca – a local discount grocery store). Retail prices (total and expressed as $/g) were obtained from the same sources or, if necessary, from another discount grocery website (superc.ca). Potential visual confounders were computed following procedures described by Blechert and colleagues[2–4]. Briefly, we quantified object size, the contributions of red, blue and green channels, contrast and brightness using customized Python scripts with scikit-image, NumPy, and PIL libraries. Minor modifications were performed relative to the original methods to consider the black background used in the present study.

When examining differences between high- and low-calorie stimuli, we found a significantly greater contribution of the blue color channel for high-calorie items ($p = 0.03$ – Fig. S1C). As expected by design, true caloric density (Fig. S2A) and displayed calories (Fig. S2C) were higher for high-calorie stimuli (both $p < 0.001$). Participants estimated caloric density was also higher for high-calorie stimuli as compared to low-calorie stimuli, indicating accurate caloric estimation ($p < 0.001$, Fig. S2B). Response times were consistently longer for high-calorie stimuli ($p < 0.05$), except at 24 months post-surgery, where the difference was not statistically significant ($p = 0.12$, Fig. S3). Pre-surgery liking ratings were lower for high-calorie stimuli ($p = 0.042$, Fig. S4A), whereas familiarity was uniformly high (near 100%) across both categories (Fig. S4B). Retail price per gram and total retail price were higher for high-calorie stimuli (both $p < 0.001$, $p = 0.012$, respectively), but the displayed weight of these stimuli was significantly lower ($p = 0.004$, Fig. S5).

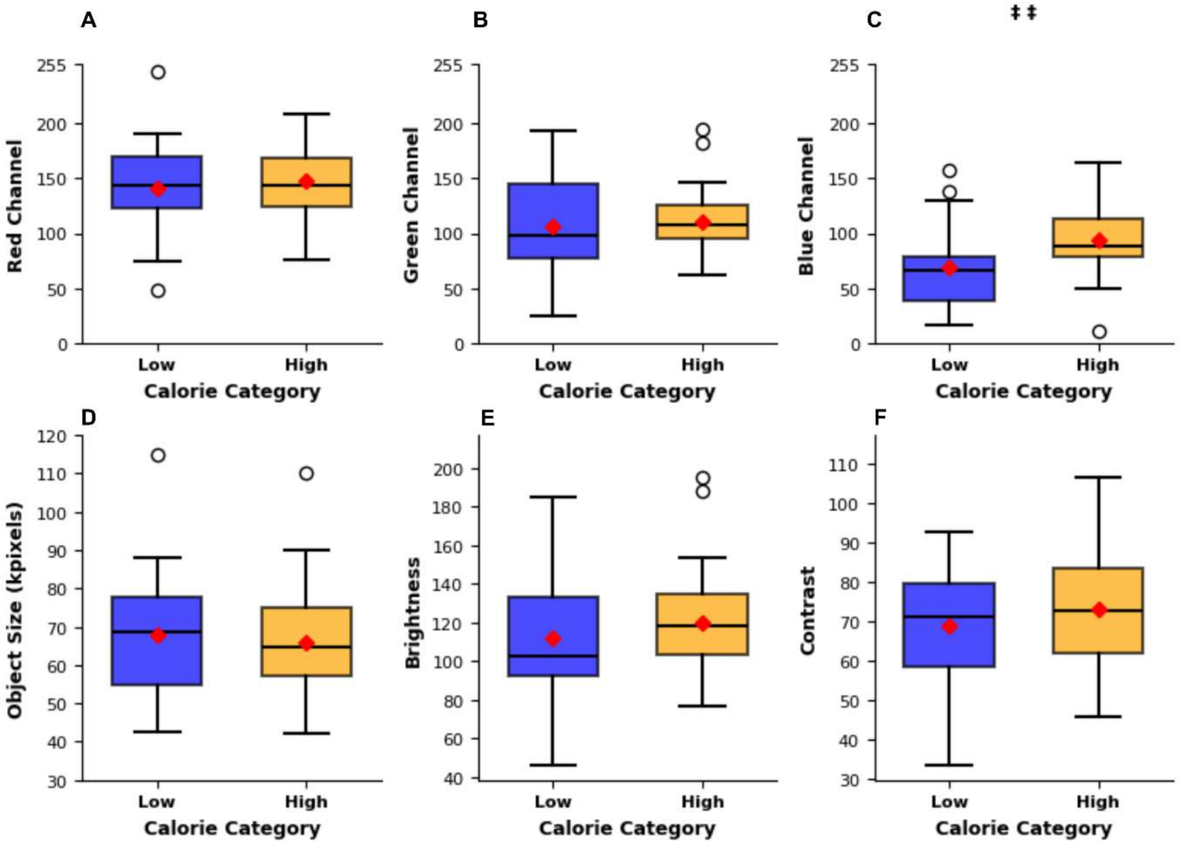


**Figure S1: Stimulus visual properties:** Standard boxplots showing the distribution of visual properties for low-calorie (blue) and high-calorie (yellow) stimuli. Red diamonds indicate the mean. ‡‡ $p<0.01$ (Mann-Whitney test)

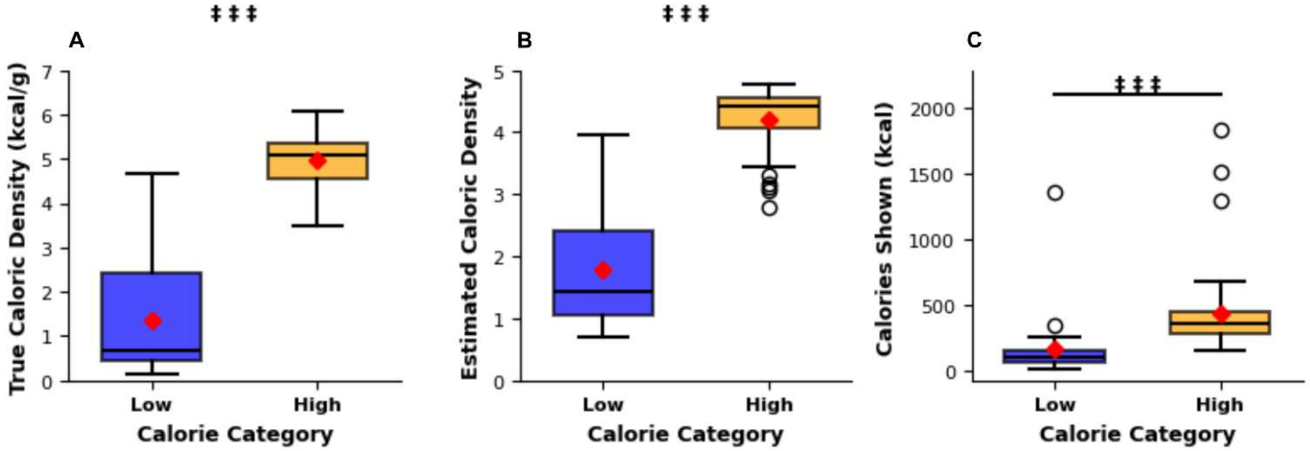


**Figure S2: Caloric properties of stimuli.** True caloric density (A), estimated caloric density at pre-surgery (B) and total displayed calories (C). Results are presented as standard boxplots showing the distribution of properties for low-calorie (blue) and high-calorie (yellow) stimuli. Red diamonds indicate the mean. ‡‡‡ $p<0.001$ (Mann-Whitney test)

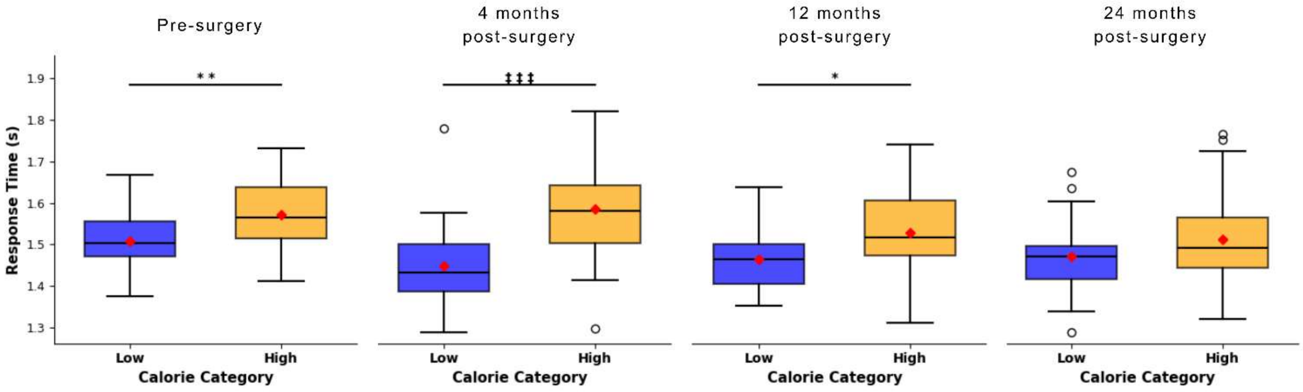


**Figure S3: Response times for stimuli across sessions.** Stimulus response times measured pre-surgery and at 4, 12 and 24 months post-surgery. Results are presented as standard boxplots showing the distribution for low-calorie (blue) and high-calorie (yellow) stimuli. Red diamonds indicate the mean. * $p<0.05$ (t-test); ** $p<0.01$ (t-test), ‡‡‡ $p<0.001$ (Mann-Whitney test)

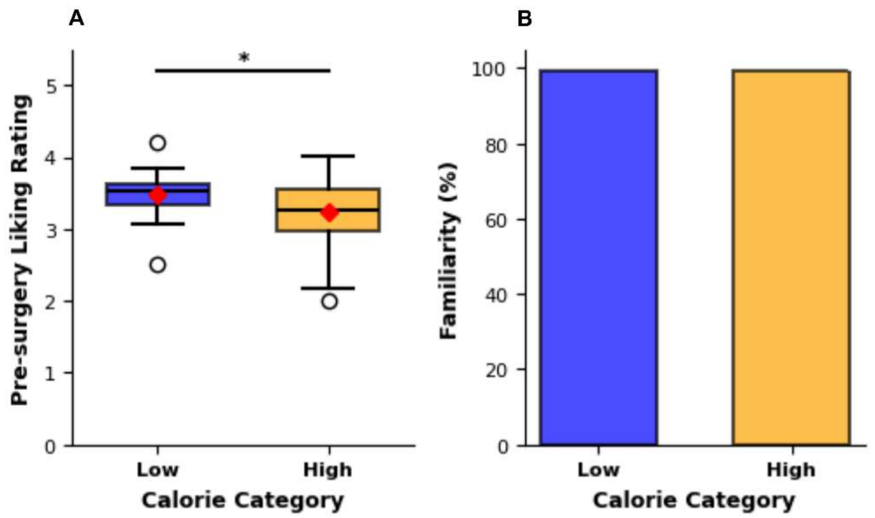


**Figure S4**: **Liking (A) and familiarity (B) ratings of stimuli at pre-surgery.** Results are presented as standard boxplots showing the distribution for low-calorie (blue) and high-calorie (yellow) stimuli. Red diamonds indicate the mean in (A). * $p<0.05$ (t-test)

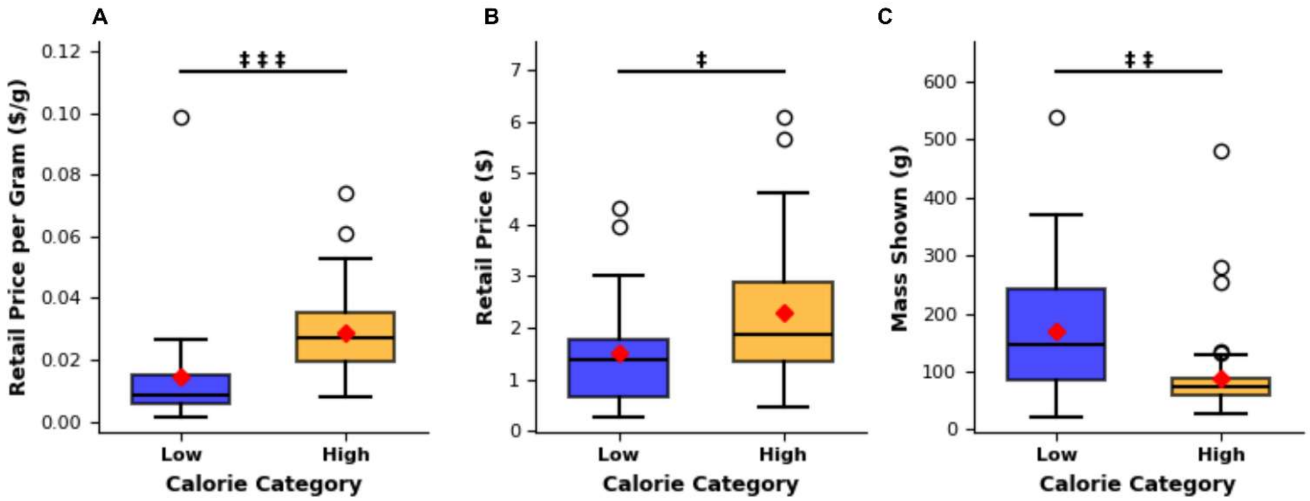


**Figure S5: Stimuli retail price per gram (A), total retail price (B) and mass shown per stimulus (C)**. Results are presented as standard boxplots showing the distribution for low-calorie (blue) and high-calorie (yellow). Red diamonds indicate the mean. ‡ $p<0.05$, ‡‡ $p<0.01$, ‡‡‡ $p<0.001$ (Mann-Whitney test)

To further examine factors influencing WTP for high- and low-calorie stimuli, additional mixed-effects models were computed. These models included TFEQ subscale scores (cognitive restraint, disinhibition, and susceptibility to hunger) at all timepoints, as well as circulating levels of fasting total ghrelin, postprandial PYY and GLP-1, as fixed effects of interest. The same covariates used in the behavioral analyses (Baseline age and BMI, surgery type, sex) were included, and participants were again modeled as a categorical random effect. For total ghrelin, PYY, and GLP-1, observations exceeding ±3 standard deviations from the session-specific group mean were excluded as outliers (n=2). The significance of parameter estimates was Bonferroni-corrected. To explore potential influences of additional stimulus properties on WTP, Pearson or Spearman correlations (depending on distributional normality) were computed between WTP amounts and the following variables: true calorie density, participants estimated calorie density (pre-surgery), pre-surgery liking ratings, and the pricing and visual properties previously identified as significantly differing between high- and low-calorie stimuli.

##### 1.2 fMRIPREP detailed preprocessing

Section 1.2 is copied in its entirety from the fMRIPREP software as requested by the authors.

Results included in this manuscript come from preprocessing performed using *fMRIPrep* 25.1.3 (Esteban et al. (2019); Esteban et al. (2018); RRID:SCR_016216), which is based on *Nipype* 1.10.0 (K. Gorgolewski et al. (2011); K. J. Gorgolewski et al. (2018); RRID:SCR_002502).

###### Preprocessing of $B_0$ inhomogeneity mappings

A total of 8 fieldmaps were found available within the input BIDS structure for this particular subject. A $B_0$ nonuniformity map (or *fieldmap*) was directly measured with an MRI scheme designed with that purpose such as SEI (Spiral-Echo Imaging).

###### Anatomical data preprocessing

A total of 4 T1-weighted (T1w) images were found within the input BIDS dataset. Each T1w image was corrected for intensity non-uniformity (INU) with N4BiasFieldCorrection (Tustison et al. 2010), distributed with ANTs 2.6.2 (Avants et al. 2008, RRID:SCR_004757). The T1w-reference was then skull-stripped with a *Nipype* implementation of the antsBrainExtraction.sh workflow (from ANTs), using OASIS30ANTs as target template. Brain tissue segmentation of cerebrospinal fluid (CSF), white-matter (WM) and gray-matter (GM) was performed on the brain-extracted T1w using fast (FSL (version unknown), RRID:SCR_002823, Zhang, Brady, and Smith 2001). An anatomical T1w-reference map was computed after registration of 4 <module 'nipype.interfaces.image' from '/opt/conda/envs/fmriprep/lib/python3.12/site-packages/nipype/interfaces/image.py'> images (after INU-correction) using mri_robust_template (FreeSurfer 7.3.2, Reuter, Rosas, and Fischl 2010). Brain surfaces were reconstructed using recon-all (FreeSurfer 7.3.2, RRID:SCR_001847, Dale, Fischl, and Sereno 1999), and the brain mask estimated previously was refined with a custom variation of the method to reconcile ANTs-derived and FreeSurfer-derived segmentations of the cortical gray-matter of Mindboggle (RRID:SCR_002438, Klein et al. 2017). Volume-based spatial normalization to one standard space (MNI152NLin2009cAsym) was performed through nonlinear registration with antsRegistration (ANTs 2.6.2), using brain-extracted versions of both T1w reference and the T1w template. The following template was selected for spatial normalization and accessed with *TemplateFlow* (24.2.2, Ciric et al. 2022): *ICBM 152 Nonlinear Asymmetrical template version 2009c* [Fonov et al. (2009), RRID:SCR_008796; TemplateFlow ID: MNI152NLin2009cAsym].

**Functional data preprocessing**

For each of the 12 BOLD runs found per subject (across all tasks and sessions), the following preprocessing was performed. First, a reference volume was generated, using a custom methodology of *fMRIPrep*, for use in head motion correction. Head-motion parameters with respect to the BOLD reference (transformation matrices, and six corresponding rotation and translation parameters) are estimated before any spatiotemporal filtering using mcflirt (FSL, Jenkinson et al. 2002). The estimated *fieldmap* was then aligned with rigid-registration to the target EPI (echo-planar imaging) reference run. The field coefficients were mapped on to the reference EPI using the transform. The BOLD reference was then co-registered to the T1w reference using bbregister (FreeSurfer) which implements boundary-based registration (Greve and Fischl 2009). Co-registration was configured with six degrees of freedom. Several confounding time-series were calculated based on the *preprocessed BOLD*: framewise displacement (FD), DVARS and three region-wise global signals. FD was computed using two formulations following Power (absolute sum of relative motions, Power et al. (2014)) and Jenkinson (relative root mean square displacement between affines, Jenkinson et al. (2002)). FD and DVARS are calculated for each functional run, both using their implementations in *Nipype* (following the definitions by Power et al. 2014). The three global signals are extracted within the CSF, the WM, and the whole-brain masks. Additionally, a set of physiological regressors were extracted to allow for component-based noise correction (*CompCor*, Behzadi et al. 2007). Principal components are estimated after high-pass filtering the *preprocessed BOLD* time-series (using a discrete cosine filter with 128s cut-off) for the two *CompCor* variants: temporal (tCompCor) and anatomical (aCompCor). tCompCor components are then calculated from the top 2% variable voxels within the brain mask. For aCompCor, three probabilistic masks (CSF, WM and combined CSF+WM) are generated in anatomical space. The implementation differs from that of Behzadi et al. in that instead of eroding the masks by 2 pixels on BOLD space, a mask of pixels that likely contain a volume fraction of GM is subtracted from the aCompCor masks. This mask is obtained by dilating a GM mask extracted from the FreeSurfer's *aseg* segmentation, and it ensures components are not extracted from voxels containing a minimal fraction of GM. Finally, these masks are resampled into BOLD space and binarized by thresholding at 0.99 (as in the original implementation). Components are also calculated separately within the WM and CSF masks. For each CompCor decomposition, the *k* components with the largest singular values are retained, such that the retained components' time series are sufficient to explain 50 percent of variance across the nuisance mask

(CSF, WM, combined, or temporal). The remaining components are dropped from consideration. The head-motion estimates calculated in the correction step were also placed within the corresponding confounds file. The confound time series derived from head motion estimates and global signals were expanded with the inclusion of temporal derivatives and quadratic terms for each (Satterthwaite et al. 2013). Frames that exceeded a threshold of 0.5 mm FD or 1.5 standardized DVARS were annotated as motion outliers. Additional nuisance timeseries are calculated by means of principal components analysis of the signal found within a thin band (*crown*) of voxels around the edge of the brain, as proposed by (Patriat, Reynolds, and Birn 2017). All resamplings can be performed with *a single interpolation step* by composing all the pertinent transformations (i.e. head-motion transform matrices, susceptibility distortion correction when available, and co-registrations to anatomical and output spaces). Gridded (volumetric) resamplings were performed using nitransforms, configured with cubic B-spline interpolation.

Many internal operations of *fMRIPrep* use *Nilearn* 0.11.1 (Abraham et al. 2014, RRID:SCR_001362), mostly within the functional processing workflow. For more details of the pipeline, see the section corresponding to workflows in *fMRIPrep*'s documentation.

###### 1.2.1 Copyright Waiver

The above boilerplate text was automatically generated by fMRIPrep with the express intention that users should copy and paste this text into their manuscripts *unchanged*. It is released under the CC0 license.

###### 1.2.2 fMRIPREP References

Abraham, Alexandre, Fabian Pedregosa, Michael Eickenberg, Philippe Gervais, Andreas Mueller, Jean Kossaifi, Alexandre Gramfort, Bertrand Thirion, and Gael Varoquaux. 2014. "Machine Learning for Neuroimaging with Scikit-Learn." *Frontiers in Neuroinformatics* 8. https://doi.org/10.3389/fninf.2014.00014.

Avants, B. B., C. L. Epstein, M. Grossman, and J. C. Gee. 2008. "Symmetric Diffeomorphic Image Registration with Cross-Correlation: Evaluating Automated Labeling of Elderly and Neurodegenerative Brain." *Medical Image Analysis* 12 (1): 26–41. https://doi.org/10.1016/j.media.2007.06.004.

Behzadi, Yashar, Khaled Restom, Joy Liau, and Thomas T. Liu. 2007. "A Component Based Noise Correction Method (CompCor) for BOLD and Perfusion Based fMRI." *NeuroImage* 37 (1): 90–101. https://doi.org/10.1016/j.neuroimage.2007.04.042.

Ciric, R., William H. Thompson, R. Lorenz, M. Goncalves, E. MacNicol, C. J. Markiewicz, Y. O. Halchenko, et al. 2022. "TemplateFlow: FAIR-Sharing of Multi-Scale, Multi-Species Brain Models." *Nature Methods* 19: 1568–71. https://doi.org/10.1038/s41592-022-01681-2.

Dale, Anders M., Bruce Fischl, and Martin I. Sereno. 1999. "Cortical Surface-Based Analysis: I. Segmentation and Surface Reconstruction." *NeuroImage* 9 (2): 179–94. https://doi.org/10.1006/nimg.1998.0395.

Esteban, Oscar, Ross Blair, Christopher J. Markiewicz, Shoshana L. Berleant, Craig Moodie, Feilong Ma, Ayse Ilkay Isik, et al. 2018. "fMRIPrep 25.1.3." *Software*. https://doi.org/10.5281/zenodo.852659.

Esteban, Oscar, Christopher Markiewicz, Ross W Blair, Craig Moodie, Ayse Ilkay Isik, Asier Erramuzpe Aliaga, James Kent, et al. 2019. "fMRIPrep: A Robust Preprocessing Pipeline for Functional MRI." *Nature Methods* 16: 111–16. https://doi.org/10.1038/s41592-018-0235-4.

Fonov, VS, AC Evans, RC McKinstry, CR Almli, and DL Collins. 2009. "Unbiased Nonlinear Average Age-Appropriate Brain Templates from Birth to Adulthood." *NeuroImage* 47, Supplement 1: S102. https://doi.org/10.1016/S1053-8119(09)70884-5.

Gorgolewski, K., C. D. Burns, C. Madison, D. Clark, Y. O. Halchenko, M. L. Waskom, and S. Ghosh. 2011. "Nipype: A Flexible, Lightweight and Extensible Neuroimaging Data Processing Framework in Python." *Frontiers in Neuroinformatics* 5: 13. https://doi.org/10.3389/fninf.2011.00013.

Gorgolewski, Krzysztof J., Oscar Esteban, Christopher J. Markiewicz, Erik Ziegler, David Gage Ellis, Michael Philipp Notter, Dorota Jarecka, et al. 2018. “Nipype.” *Software*. https://doi.org/10.5281/zenodo.596855.

Greve, Douglas N, and Bruce Fischl. 2009. “Accurate and Robust Brain Image Alignment Using Boundary-Based Registration.” *NeuroImage* 48 (1): 63–72. https://doi.org/10.1016/j.neuroimage.2009.06.060.

Jenkinson, Mark, Peter Bannister, Michael Brady, and Stephen Smith. 2002. “Improved Optimization for the Robust and Accurate Linear Registration and Motion Correction of Brain Images.” *NeuroImage* 17 (2): 825–41. https://doi.org/10.1006/nimg.2002.1132.

Klein, Arno, Satrajit S. Ghosh, Forrest S. Bao, Joachim Giard, Yrjö Häme, Eliezer Stavsky, Noah Lee, et al. 2017. “Mindboggling Morphometry of Human Brains.” *PLOS Computational Biology* 13 (2): e1005350. https://doi.org/10.1371/journal.pcbi.1005350.

Patriat, Rémi, Richard C. Reynolds, and Rasmus M. Birn. 2017. “An Improved Model of Motion-Related Signal Changes in fMRI.” *NeuroImage* 144, Part A (January): 74–82. https://doi.org/10.1016/j.neuroimage.2016.08.051.

Power, Jonathan D., Anish Mitra, Timothy O. Laumann, Abraham Z. Snyder, Bradley L. Schlaggar, and Steven E. Petersen. 2014. “Methods to Detect, Characterize, and Remove Motion Artifact in Resting State fMRI.” *NeuroImage* 84 (Supplement C): 320–41. https://doi.org/10.1016/j.neuroimage.2013.08.048.

Reuter, Martin, Herminia Diana Rosas, and Bruce Fischl. 2010. “Highly Accurate Inverse Consistent Registration: A Robust Approach.” *NeuroImage* 53 (4): 1181–96. https://doi.org/10.1016/j.neuroimage.2010.07.020.

Satterthwaite, Theodore D., Mark A. Elliott, Raphael T. Gerraty, Kosha Ruparel, James Loughead, Monica E. Calkins, Simon B. Eickhoff, et al. 2013. “An improved framework for confound regression and filtering for control of motion artifact in the preprocessing of resting-state functional connectivity data.” *NeuroImage* 64 (1): 240–56. https://doi.org/10.1016/j.neuroimage.2012.08.052.

Tustison, N. J., B. B. Avants, P. A. Cook, Y. Zheng, A. Egan, P. A. Yushkevich, and J. C. Gee. 2010. “N4ITK: Improved N3 Bias Correction.” *IEEE Transactions on Medical Imaging* 29 (6): 1310–20. https://doi.org/10.1109/TMI.2010.2046908.

Zhang, Y., M. Brady, and S. Smith. 2001. “Segmentation of Brain MR Images Through a Hidden Markov Random Field Model and the Expectation-Maximization Algorithm.” *IEEE Transactions on Medical Imaging* 20 (1): 45–57. https://doi.org/10.1109/42.906424.

##### 1.3 Quality control

Quality control was informed using MRIQC-derived metrics[5], with runs flagged when indicators suggested poor data quality. Runs with a maximum framewise displacement (FDmax) exceeding 3 mm were flagged. Because the ventromedial prefrontal cortex (vmPFC) was a region of interest (ROI) and is particularly susceptible to signal loss, vmPFC coverage was evaluated for each functional brain mask by comparison with a vmPFC mask derived from the BDM subjective valuation meta-analysis by Newton-Fenner et al.[6]. Runs with less than 90% coverage were excluded. Additional quality control flags were applied to runs in which at least one metric exceeded 1 z-score above the session-wide mean for framewise displacement [7], AFNI’s quality index (AQI)[8], and AFNI’s AOR score [8], or fell more than 1 z-score below the mean for total signal-to-noise ratio[9]. Two independent reviewers (CL and PG) visually inspected fMRIPREP reports. Exclusion criteria were defined based on previously recommended procedures[10] and included excessive motion artefacts, important signal loss, periodicity and preprocessing failures.

##### 1.4 Details on participant-level modeling

Using Nilearn[11], preprocessed functional data were spatially smoothed using an 8 mm full-width at half-maximum (FWHM) Gaussian kernel, scaled to voxelwise percent signal change, and high-pass filtered using 9 discrete cosine drift regressors. Nuisance regressors included six motion parameters (three translational and three rotational) and signals extracted from cerebrospinal fluid (CSF) and white matter (WM) masks. Temporal autocorrelation was modeled using a first-order autoregressive (AR1) model. For the high- versus low-calorie contrast, whether modulated or unmodulated, the contrast vector was specified as [0.5, 0.5, −1] for high-calorie savory, high-calorie sweet, and low-calorie stimuli, respectively, with all remaining regressors assigned a weight of zero. This approach allowed for a balanced comparison between high- and low-calorie food categories by averaging across high-calorie sweet and savory subtypes.

##### 1.5 Details on second-level modeling

In both group-level models (Model I and II), categorical variables included session (pre-surgery, 4, 12 and 24 months post-surgery), surgery type (SG, RYGB, BPD-DS), handedness (right or left) and run (1, 2 or 3). Baseline BMI, baseline age and total weight loss (TWL) at 24 months were mean-centered. Hunger ratings collected immediately before each scan were extracted at each session and grand-mean centered across all participants and timepoints. A 50% thresholded brain mask was computed from all participant-level maps to constrain analyses to voxels present in the majority of participants' scans[8]. To estimate noise, residuals smoothness was computed using AFNI's 3dFWHMx on the residual maps generated by the first-level models, providing the ACF noise spatial smoothness parameters (≈13 mm) required for cluster-level correction.

##### 1.6 Participant inclusion workflow

A total of 89 participants were enrolled in the full study. Thirteen were excluded prior to analysis due to incomplete imaging data, resulting from study withdrawal (n = 7), technical issues during MRI acquisition (n = 1), medical complications (n = 2), or surgery-related logistical constraints (n = 3). This yielded 76 participants whose data proceeded to quality control.

Across all these participants, 779 fMRI runs were available. Quality control procedures resulted in the removal of 146 runs due to excessive motion, 30 due to signal loss, 13 due to periodicity artifacts, and 2 due to preprocessing failures. Following this step, an additional 19 participants were excluded because they lacked usable pre-surgery fMRI data or had fewer than two valid imaging sessions remaining.

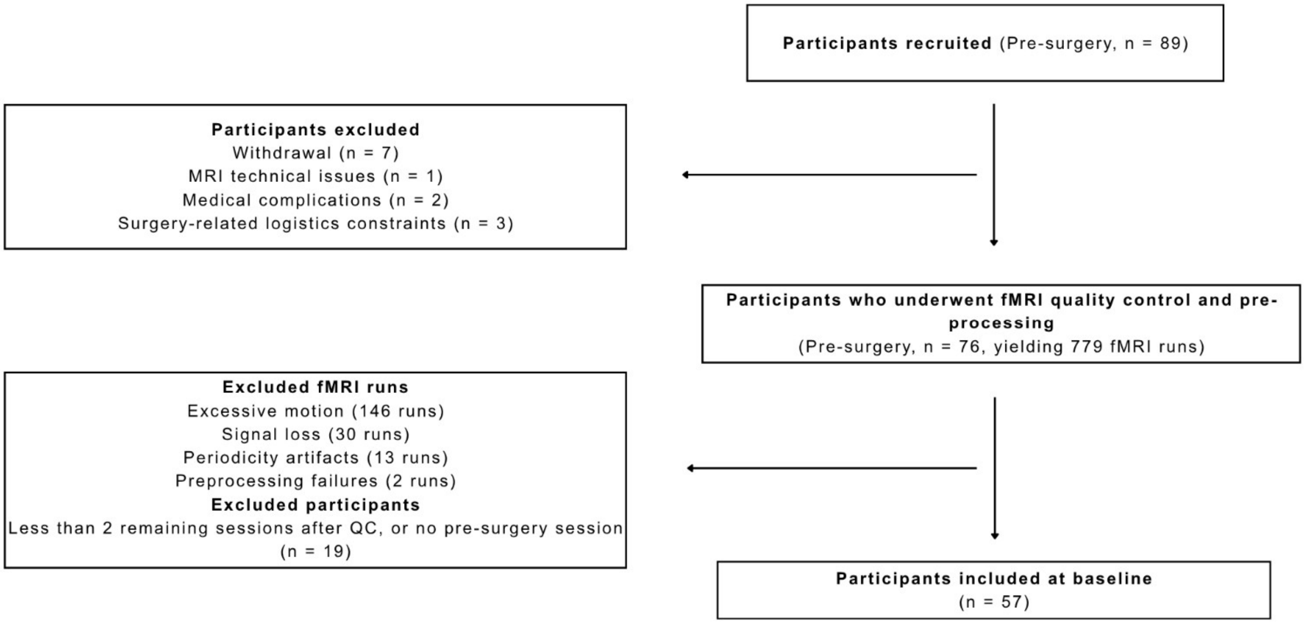


**Figure S6: Flowchart of participants included in the study**

#### 2 Supplementary results

##### 2.1 Supplementary results of willingness-to-pay and stimulus properties

To replicate previous findings[12–14], we also examined associations between WTP and true caloric density (Fig. S7), estimated caloric density (Fig. S8), and liking ratings (Fig. S9). Before surgery, WTP for all stimuli combined was not correlated with true caloric density ($r_s = 0.11$, $p = 0.363$). However, after surgery, this association became significantly negative (all $r_s < -0.28$, all $p < 0.02$, Fig. S7). When examining stimulus categories separately, WTP for high-calorie stimuli was positively correlated with true caloric density at all sessions (all $r_s > 0.33$, all $p < 0.021$), whereas no correlations were observed for low-calorie stimuli (all $p > 0.05$, Fig. S7). Before surgery, WTP for all stimuli combined was not correlated with estimated caloric density ($r_s = 0.03$, $p = 0.824$), but after surgery, this association became significantly negative (all $r_s < -0.57$, all $p < 0.001$, Fig. S8). When analyzing categories independently, WTP for high-calorie stimuli was positively correlated with estimated caloric density pre-surgery ($r_s = 0.40$, $p = 0.005$), but after surgery, this relationship became significantly negative at all timepoints (all $r_s < -0.34$, all $p < 0.016$) except at 24 months ($r_s = -0.21$, $p = 0.146$, Fig. S8). Again, no correlations were found for low-calorie stimuli ($p > 0.216$). At all timepoints, WTP was positively correlated with pre-surgery liking ratings for both stimulus categories (all $r_s > 0.32$, all $p < 0.006$), but not at 4 months ($r_s = -0.01$, $p = 0.933$) and 12 months ($r_s = 0.27$, $p = 0.062$) post-surgery, where no significant correlation was found for high-calorie stimuli (Fig. S9).

Because price may be a potential confound, we examined associations between WTP and price per gram (Fig. S10) as well as total retail price (Fig. S11). Although no significant correlations were observed pre-surgery, price per gram was negatively and significantly correlated with WTP for all stimuli combined after surgery (all $r_s < -0.28$, all $p < 0.021$). No significant effects were observed when examining stimulus categories separately (all $p > 0.05$). A similar pattern was observed for total retail price; however, these correlations did not reach statistical significance.

Response time was negatively correlated with WTP at 4 ($r_s = -0.72$, $p < 0.001$) and 12 months post-surgery ($r_s = -0.24$, $p = 0.046$), but no significant correlations were found pre-surgery ($r_s = 0.22$, $p = 0.065$) or at 24 months post-surgery ($r_s = -0.19$, $p = 0.11$, Fig. S13). This effect appeared to be driven by high-calorie stimuli at 4 months post-surgery ($r_s = -0.51$, $p < 0.001$).

We also assessed correlations with blue-channel intensity, which significantly differed between high- and low-calorie stimuli (Fig. S14). WTP was negatively correlated with mean blue-channel intensity, but only when considering all stimuli combined, and only at 4 ($r_s = -0.26$, $p < 0.031$) and 12 months post-surgery ($r_s = -0.25$, $p = 0.037$, Fig. S14).

##### 2.2 Supplementary Figures and Tables

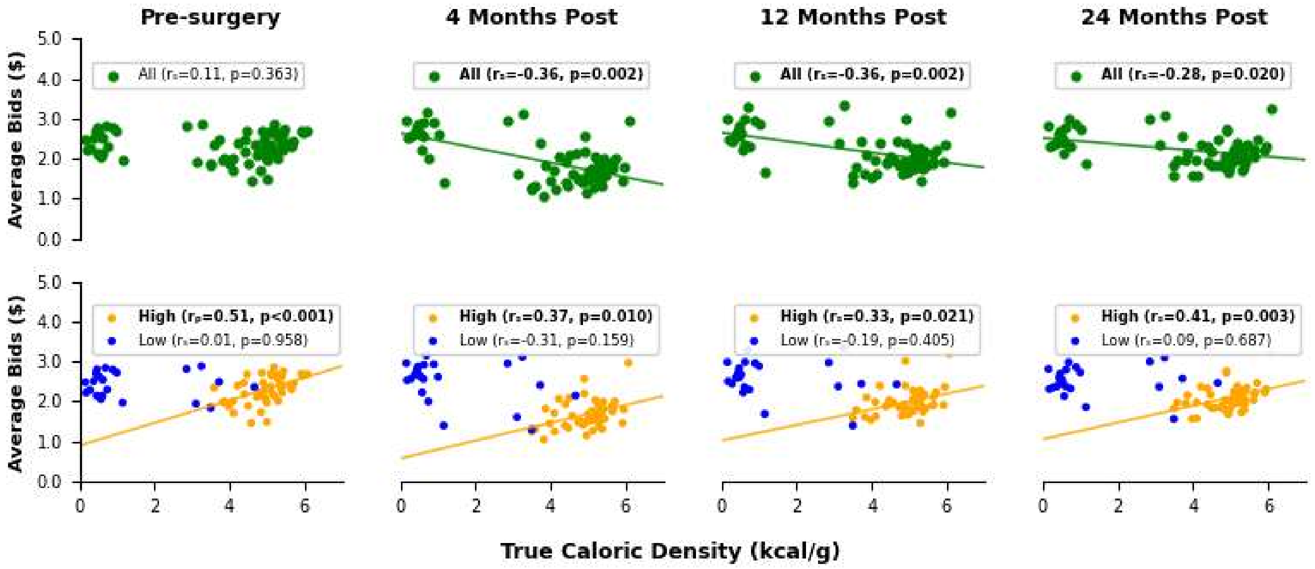


**Figure S7: Associations between true caloric density and willingness-to-pay (WTP) across timepoints.** Top panels show correlations for all stimuli combined; whereas bottom panels show correlations stratified by caloric category (yellow = high-calorie; blue = low-calorie). Pearson's correlation coefficient ($r_p$) was used when normality assumptions were met (Shapiro-Wilk $p > 0.05$); otherwise, Spearman's correlation coefficient ($r_s$) was used. Regression lines shown when *r* coefficient $p < 0.05$. Bold text also indicates statistically significant correlations ($p < 0.05$).

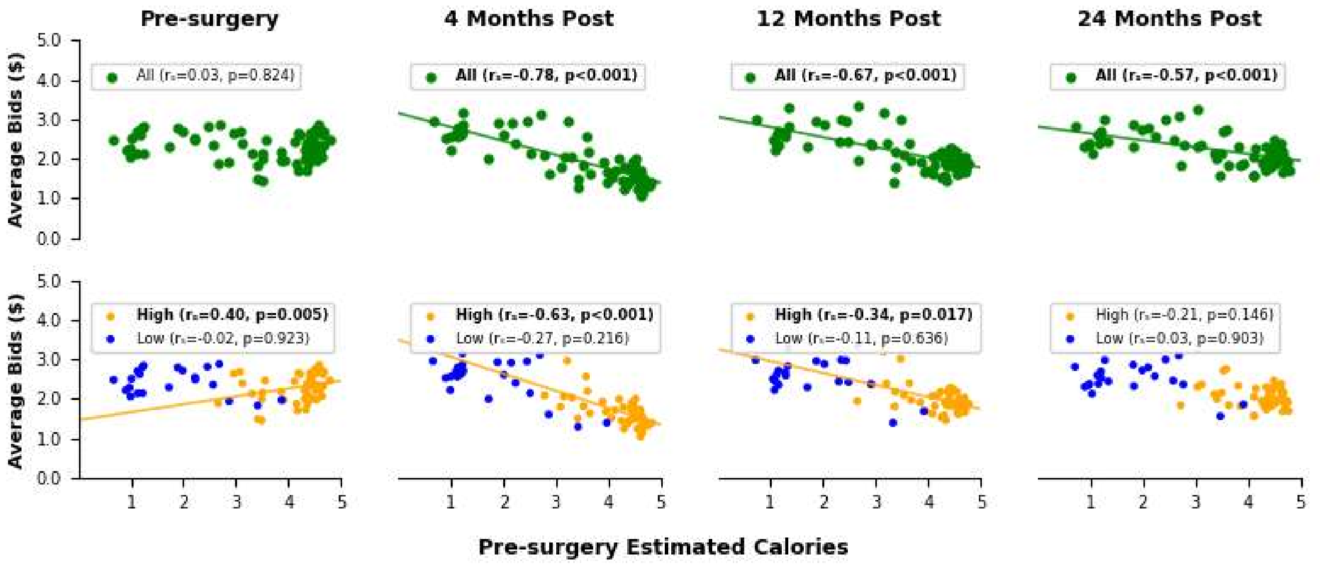


**Figure S8: Associations between pre-surgery estimated caloric density and willingness-to-pay (WTP) across timepoints.** Top panels show correlations for all stimuli combined; whereas bottom panels show correlations stratified by caloric category (yellow = high-calorie; blue = low-calorie). Pearson's correlation coefficient ($r_p$) was used when normality assumptions were met (Shapiro-Wilk p > 0.05); otherwise, Spearman's correlation coefficient ($r_s$) was used. Regression lines shown when *r* coefficient $p < 0.05$. Bold text also indicates statistically significant correlations ($p < 0.05$).

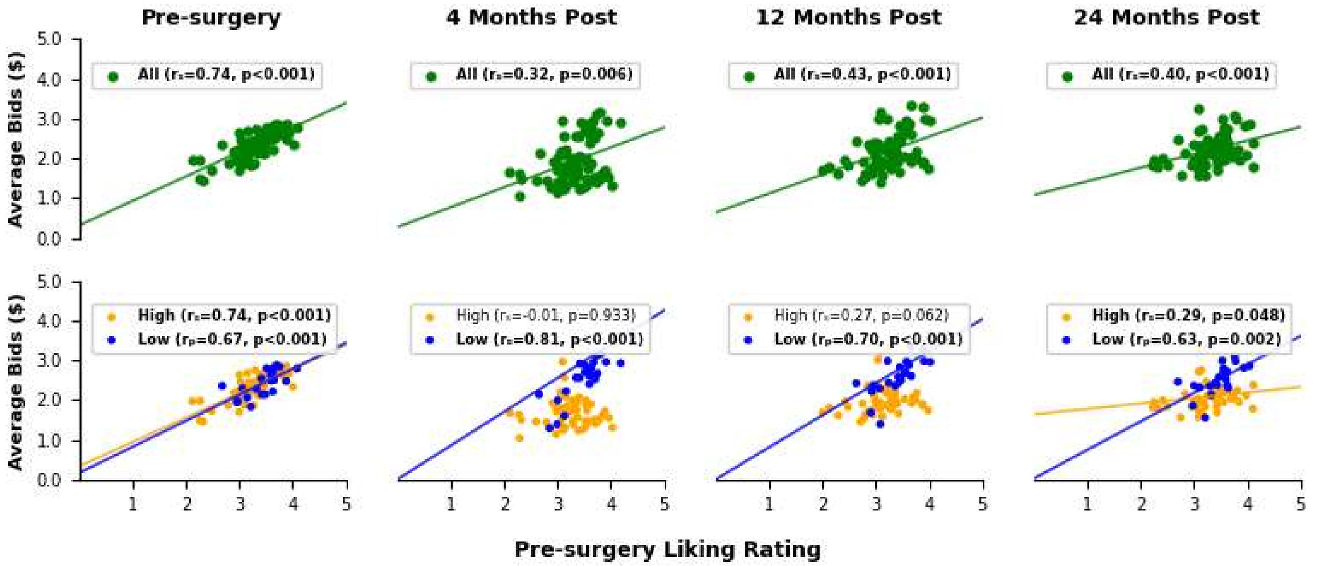

**Figure S9: Associations between pre-surgery liking ratings and willingness-to-pay (WTP) across timepoints.** Top panels show correlations for all stimuli combined; whereas bottom panels show correlations stratified by caloric category (yellow = high-calorie; blue = low-calorie). Pearson's correlation coefficient ($r_p$) was used when normality assumptions were met (Shapiro-Wilk $p > 0.05$); otherwise, Spearman's correlation coefficient ($r_s$) was used. Regression lines shown when *r* coefficient $p < 0.05$. Bold text also indicates statistically significant correlations ($p < 0.05$).

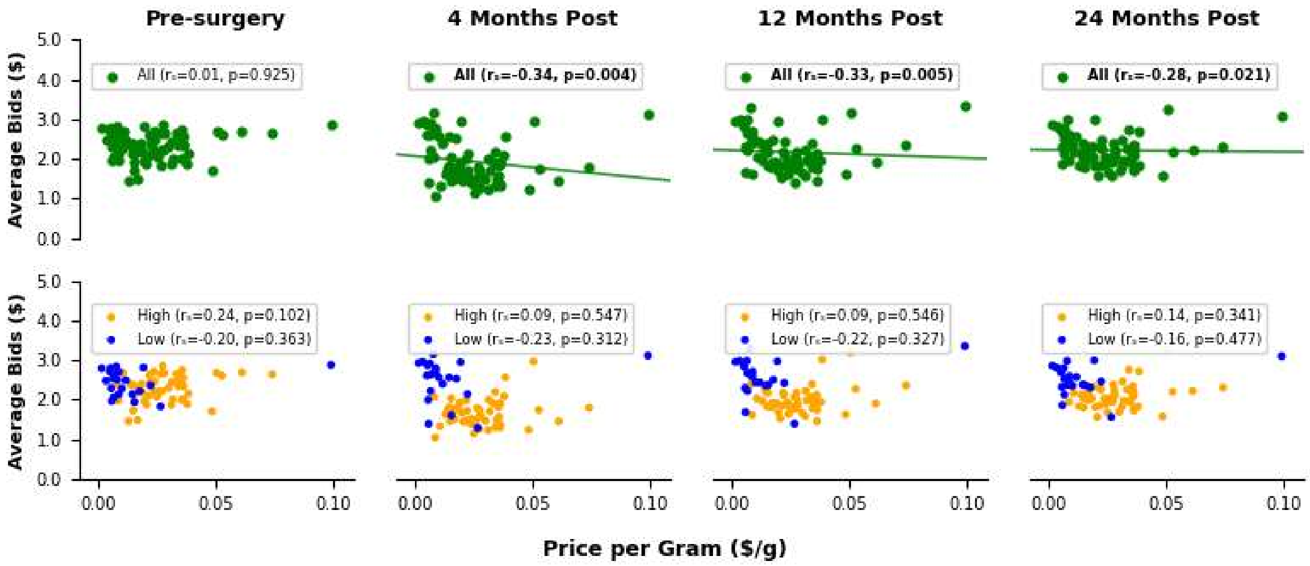


**Figure S10: Associations between price per gram and willingness-to-pay (WTP) across timepoints.** Top panels show correlations for all stimuli combined; whereas bottom panels show correlations stratified by caloric category (yellow = high-calorie; blue = low-calorie). Pearson's correlation coefficient ($r_p$) was used when normality assumptions were met (Shapiro-Wilk $p > 0.05$); otherwise, Spearman's correlation coefficient ($r_s$) was used. Regression lines shown when *r* coefficient $p < 0.05$. Bold text also indicates statistically significant correlations ($p < 0.05$).

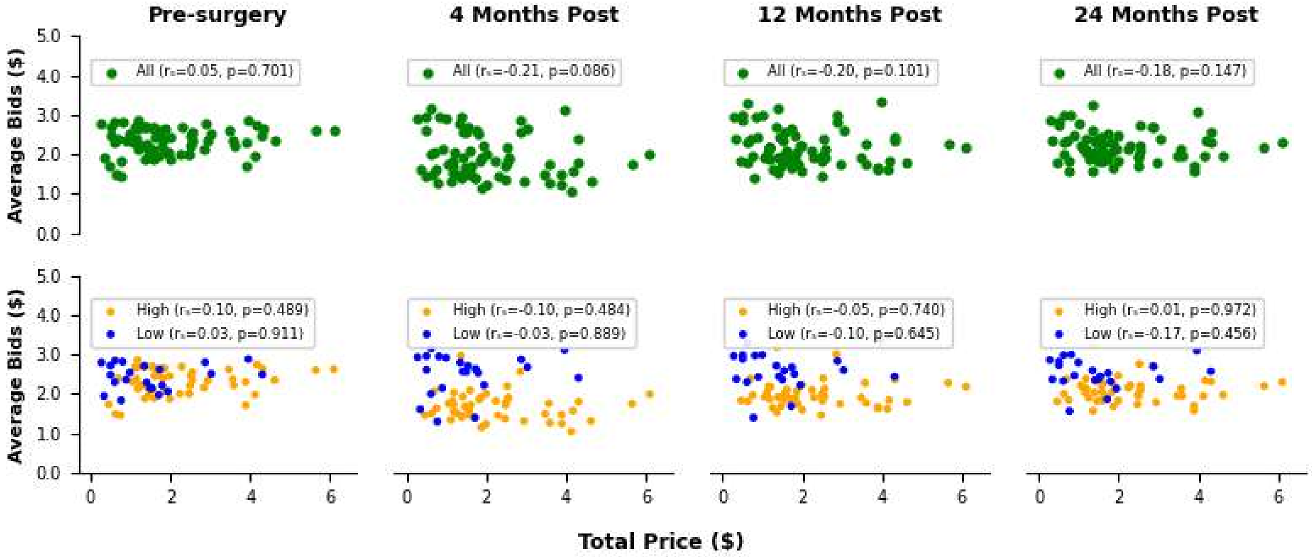


**Figure S11: Associations between total price and willingness-to-pay (WTP) across timepoints.** Top panels show correlations for all stimuli combined; whereas bottom panels show correlations stratified by caloric category (yellow = high-calorie; blue = low-calorie). Pearson's correlation coefficient ($r_p$) was used when normality assumptions were met (Shapiro-Wilk p > 0.05); otherwise, Spearman's correlation coefficient ($r_s$) was used. Regression lines shown when *r* coefficient p < 0.05. Bold text also indicates statistically significant correlations (p < 0.05).

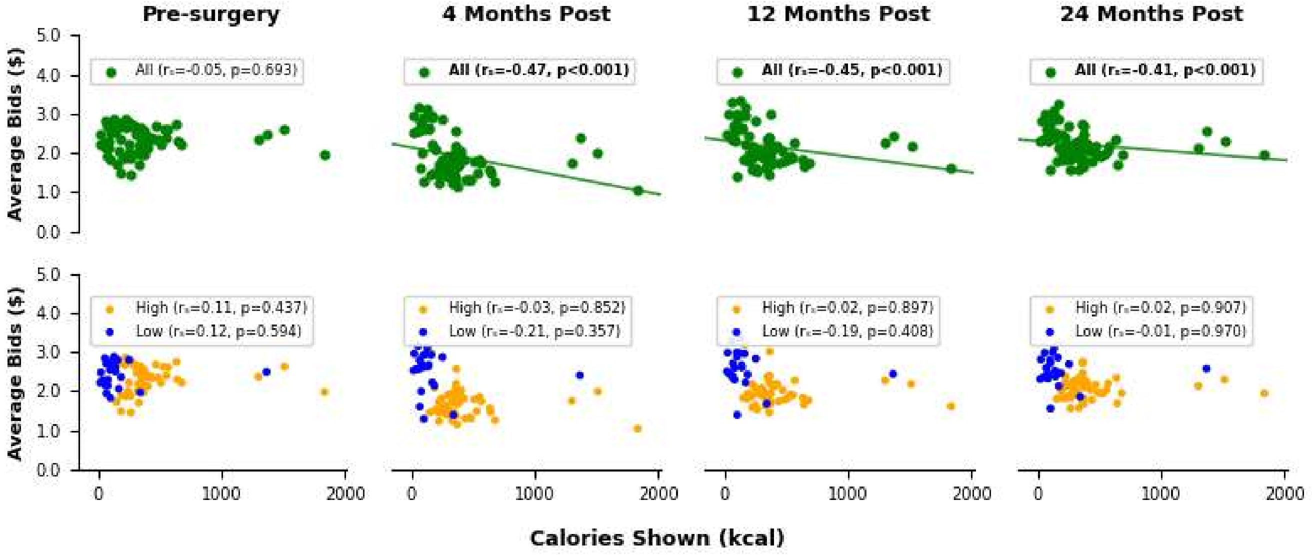

**Figure S12: Associations between total calories shown and willingness-to-pay (WTP) across timepoints.** Top panels show correlations for all stimuli combined; whereas bottom panels show correlations stratified by caloric content (yellow = high-calorie; blue = low-calorie). Pearson's correlation coefficient ($r_p$) was used when normality assumptions were met (Shapiro-Wilk $p > 0.05$); otherwise, Spearman's correlation coefficient ($r_s$) was used. Regression lines shown when *r* coefficient $p < 0.05$. Bold text also indicates statistically significant correlations ($p < 0.05$).

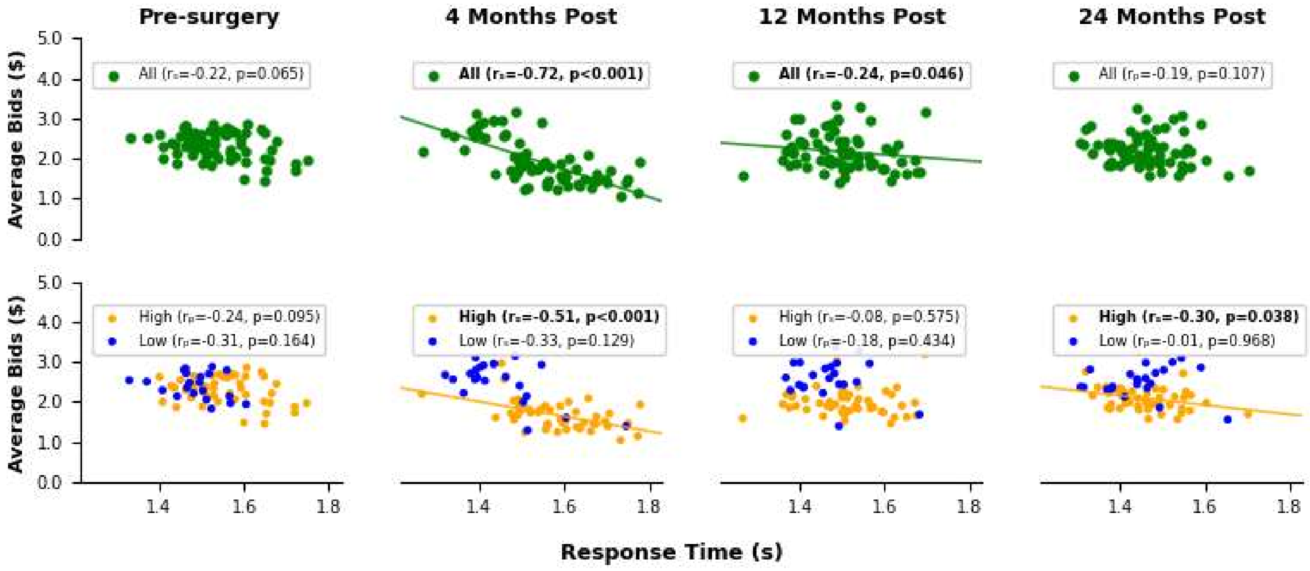


**Figure S13: Associations between response time and willingness-to-pay (WTP) across timepoints.** Top panels show correlations for all stimuli combined; whereas bottom panels show correlations stratified by calorie category (yellow = high-calorie; blue = low-calorie). Pearson's correlation coefficient ($r_p$) was used when normality assumptions were met (Shapiro-Wilk $p > 0.05$); otherwise, Spearman's correlation coefficient ($r_s$) was used. Regression lines shown when *r* coefficient $p < 0.05$. Bold text also indicates statistically significant correlations ($p < 0.05$).

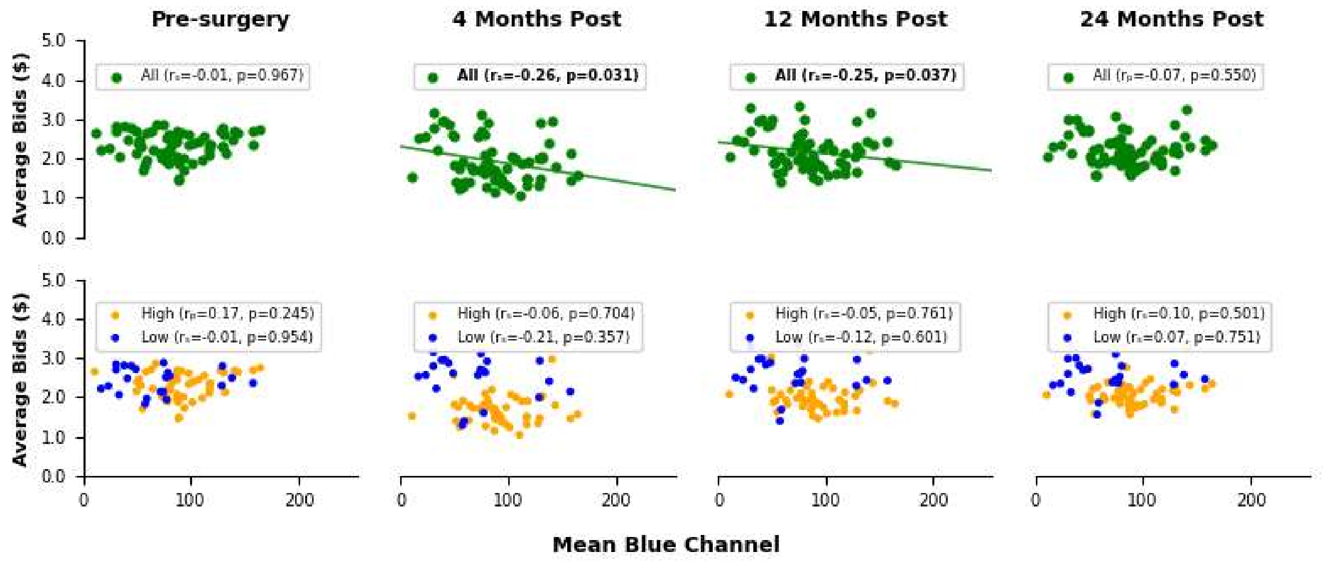


**Figure S14: Association between mean blue-channel intensity and willingness-to-pay across bariatric surgery timepoints.** Top panels show correlations for all stimuli combined; whereas bottom panels show correlations stratified by calorie category (yellow = high-calorie; blue = low-calorie). Pearson's correlation coefficient ($r_p$) was used when normality assumptions were met (Shapiro-Wilk p > 0.05); otherwise, Spearman's correlation coefficient ($r_s$) was used. Regression lines shown when *r* coefficient $p < 0.05$. Bold text also indicates statistically significant correlations ($p < 0.05$).

##### 2.2.1 Table S1: Clusters for whole-brain results - covariates

| Model | Group-level effect or contrast | Participant-level effect or contrast | Cluster size | Peak Z | Peak MNI | Mean ES | Brainnetome label and name | Coverage % |
|---|---|---|---|---|---|---|---|---|
| Model I (Session) | Surgery type (BPD > SG) | WTP-BOLD association high-calorie > low-calorie | 55 | 4.28 | -48, 36, -18 | -0.30[a] | Left orbital gyrus orbital area 12/47 | 36.4 |
| | | | | | | | Left orbital gyrus lateral area 12/47 | 32.6 |
| | | | 37 | 4.23 | -63, 12, 33 | -0.18[a] | Area 4 (head and face region) | 1.5 |
| | Run 2 > Run 1 | WTP-BOLD association high-calorie > low-calorie | 42 | 4.73 | -30, 69, -3 | 0.21[a] | Left middle frontal gyrus lateral area 10 | 9.1 |

| Model | Group-level effect or contrast | Participant-level effect or contrast | Cluster size | Peak Z | Peak MNI | Mean ES | Brainnetome label and name | Coverage % |
|---|---|---|---|---|---|---|---|---|
| | Run 3 > Run 1 | WTP-BOLD association high-calorie > low-calorie | 51 | 4.22 | 30, 45, -15 | 0.17[a] | Right orbital gyrus lateral area 11 | 80.5 |
| | | | 39 | 4.21 | 60, 3, 24 | 0.09[a] | Right precentral gyrus area 4 (head and face region) | 53.9 |
| | Handedness (left > right) | WTP-BOLD association high-calorie > low-calorie | 43 | 4.04 | -12, 12, 33 | 0.14[a] | Left cingulate gyrus caudodorsal area 24 | 18.6 |
| | | | 42 | 4.74 | -30, -12, -30 | -0.44[a] | Left parahippocampus rostral area 35/36 | 38.5 |
| | | | | | | | Left rostral hippocampus | 32.5 |
| | | Viewing high-calorie > low-calorie | 66 | 4.08 | 57, -21, 36 | 0.14[b] | Right inferior parietal rostrodorsal area 40(PFt) | 58.8 |
| | | | | | | | Right Postcentral gyrus area 2 | 23.5 |
| Model II (SessionXTWL) | Surgery type (RYGB > SG) | Viewing high-calorie > low-calorie | 35 | 3.78 | -3, 18, 30 | -0.12[b] | Right cingulate gyrus pregenual area 32 | 26.2 |
| | | | | | | | Left cingulate gyrus caudodorsal area 24 | 24.4 |
| | Surgery type (BPD > SG) | WTP-BOLD association high-calorie > low-calorie | 37 | 3.84 | -45, 42, -18 | -0.30[a] | Left orbital area 12/47 | 41.6 |
| | | | | | | | Left middle frontal gyrus lateral area10 | 32.7 |
| | Run 3 > Run 1 | WTP-BOLD association high-calorie > low-calorie | 66 | 3.90 | 39, -30, 54 | 0.08[a] | Right Postcentral gyrus area 2 | 40.2 |
| | | | | | | | Right Precentral gyrus area 4(upper limb region) | 23.6 |
| | | | 59 | 4.72 | 66, 15, 24 | 0.10[a] | Right Precentral gyrus area 4 (head and face region) | 39.2 |
| | Handedness (left > right) | WTP-BOLD association | 35 | 4.10 | -12, 6, 36 | 0.16[a] | Right Cingulate gyrus caudodorsal area 24 | 16.7 |

| Model | Group-level effect or contrast | Participant-level effect or contrast | Cluster size | Peak Z | Peak MNI | Mean ES | Brainnetome label and name | Coverage % |
|---|---|---|---|---|---|---|---|---|
| | | high-calorie > low-calorie | | | | | | |
| | | Viewing high-calorie > low-calorie | 61 | 4.13 | 57, -21, 36 | 0.16[b] | Right Inferior parietal rostrodorsal area 40(PFt) | 52.2 |
| | | | | | | | Right Postcentral gyrus area 2 | 24.1 |

**Table legend**: Covariate results from linear mixed-effects models examining brain activation during food valuation tasks. Two models were tested: Model I assessed session effects with covariates (age, sex, baseline BMI, surgery type, run and hunger before scan), and Model II assessed the interaction between session and total weight loss (TWL) at 24 months, including the same covariates. Clusters were identified using a voxel-wise threshold of $p < 0.001$ and corrected for multiple comparisons using noise autocorrelation function (ACF) estimation at FWE = 0.05. Regions with ≥ 20% coverage are reported; otherwise, the principal (first) region of the cluster is reported. Column definitions are as follows: Model: Statistical model specification, Second-Level Contrast: covariate or categorical contrast, Participant-Level Contrast: Task-related activation contrast, Cluster Size: Number of voxels in the cluster. Peak Z: Maximum Z-score within the cluster, Peak MNI: MNI coordinates (x, y, z) of the peak activation in mm, Mean ES: Mean effect size within the cluster, expressed as (a) BOLD percent signal change per $ relative to the run mean, per categorical contrast (e.g. BPD-SG vs. SG), or (b) BOLD percent signal change per categorical contrast. Brainnetome Label: Anatomical region from Brainnetome Atlas[15], Coverage %: Percentage of the cluster overlapping with labeled region. Regions with ≥ 20% overlap are reported, unless all overlaps are < 20%, in which case the region with the greatest overlap is shown. Abbreviations: TWL, total weight loss; ES, effect size; MNI, Montreal Neurological Institute coordinate space, BPD: biliopancreatic diversion with duodenal switch, RYGB: Roux-en-Y gastric bypass, SG: Sleeve gastrectomy.
